# Bait, not reward: CO_2_-enriched *Nepenthes* pitchers secrete toxic nectar

**DOI:** 10.1101/2023.11.25.568661

**Authors:** Chandni Chandran Lathika, Gokul Baburaj Sujatha, Gintu Thomas, Anil John Johnson, Gayathri Viswanathan, Thania Sara Varghese, Saleem Mohamed, Lubaina Abdulhadeef Shereefa, Sabulal Baby

## Abstract

*Nepenthes* pitchers are leaf-evolved biological traps holding high levels of CO_2_ within them. Extrafloral nectar (EFN) secreted by these pitchers has long been regarded as the major reward to visiting arthropods, but its chemical constituents and their role in prey capture are least explored. Here we demonstrate *Nepenthes* EFN as a sugar (glucose-fructose-sucrose) mix with high C:N ratio, minimal amino acids, proteins, and vitamin C. *Nepenthes khasiana* peristome and lid EFNs displayed strong acetylcholinesterase (AChE) inhibition; the naphthoquinone derivative, (+)-isoshinanolone, has been identified as the AChE inhibitor. Plumbagin, the major volatile naphthoquinone in *Nepenthes*, also showed strong AChE inhibition. Direct EFN– and (+)-isoshinanolone-feeding bioassays demonstrated symptoms of cholinergic toxicity in ants. We testify that *Nepenthes* EFN is a toxic bait which hinders neuronal activity in visiting arthropods. These unique traps adopt various deceptive strategies for prey capture, and our discovery abolishes the notion that *Nepenthes* EFN is a reward to visiting ants and other arthropods. Moreover, our findings infer elevated CO_2_ within their pitchers as the key factor influencing the growth, metabolism, herbivory, and carnivory in *Nepenthes*.

**Highlight:** *Nepenthes* extrafloral nectar has high content of carbohydrates and minimal nitrogenous metabolites. It is laced with (+)-isoshinanolone, an acetylcholinesterase inhibitor, and acts as a toxic bait, aiding prey capture.

## Introduction

*Nepenthes* pitchers effect prey capture and nutrient acquisition through various physico-chemical and biological strategies (Bohn and Federle, 2004; Baby *et al*., 2017; Mithöfer, 2022). Their pitcher rim (peristome) is a wettable, low-friction surface with distinct microstructures (Bohn and Federle, 2004; Chen *et al*., 2016), and these are crucial factors deciding the efficiency of prey capture. Environmental factors (humidity, rain) also influence the wettability of prey trapping surface and capture rates (Bohn and Federle, 2004; Chen *et al*., 2016; Sujatha *et al*., 2023). In addition, visual signals were reported from prey capturing regions of *Nepenthes* pitchers (Joel *et al*., 1985; Moran *et al*., 1999; Kurup *et al*., 2013). We have described *Nepenthes* pitchers as CO_2_-enriched cavities, and open pitchers continually emit CO_2_ attracting preys (Baby *et al*., 2017). Over the years researchers reported the role of pitcher colour (Bennett and Ellison, 2009; Dávila-Lara *et al*., 2021a), scent (Di Giusto *et al*., 2008; Jürgens *et al*., 2009) and peristome geometry (Moulton *et al*., 2023) in prey capture in *Nepenthes*.

Extrafloral nectar (EFN) produced by nectaries on the tendrils, peristomes and lids of *Nepenthes* pitchers is considered as the primary nutritional reward to the visiting ants, other insects and small mammals (Bauer *et al*., 2008; Bennett and Ellison, 2009; Gaume *et al*., 2016). EFN also contributes to peristome wettability (Bauer *et al*., 2008; Sujatha *et al*., 2023). Bornean montane *Nepenthes* species through mutualistic interactions provide nectar and harvest nitrogenous excreta from tree shrews and other small mammals with their shape-adapted pitchers (Greenwood *et al*., 2011). Non-mutualistic interactions of diverse vertebrates with *Nepenthes* pitchers as their nectar (food) source were also reported from Brunei (Bauer *et al*., 2016). In angiosperms, floral nectar (FN) is one of the major rewards in pollination (Nicolson, 2022). Nectar, being a sweet aqueous secretion, is easily consumed, digested and absorbed. It has a complex chemistry containing sugars, amino acids, proteins, phenolics, alkaloids, lipids, organic acids and inorganic ions (De la Barrera and Nobel, 2004; Nepi *et al*., 2012; Nicolson, 2022). However, the composition of nectar produced by floral and extrafloral nectaries in *Nepenthes* is barely investigated (Heil, 2015; Hatcher *et al*., 2020). In this study, we describe the role of EFN produced by *Nepenthes* pitchers in prey capture by deciphering its chemistry, prey spectrum and insect toxicity. The results are interpreted in the backdrop of the discovery of high levels of CO_2_ within *Nepenthes* pitchers (Baby *et al*., 2017).

## Materials and methods

### *Nepenthes* species, hybrids

*Nepenthes khasiana* Hook.f., *N. mirabilis* (Lour.) Druce and hybrids *N. mirabilis* (Lour.) Druce × *N. khasiana* Hook.f., *N. mirabilis* (Lour.) Druce × *N. rafflesiana* Jack pitchers were collected from the conservatories of Jawaharlal Nehru Tropical Botanic Garden and Research Institute (JNTBGRI) (08° 45′00.04″N, 77° 01′41.09″E) from August 2023 to November 2024. Voucher specimens of *Nepenthes* species/hybrids (*N. khasiana* TBGT 39201, *N. mirabilis* TBGT 44774, *N. mirabilis × N. khasiana* TBGT 44776, *N. mirabilis × N. rafflesiana* TBGT 44777) were deposited at JNTBGRI Herbarium.

### EFN isolation

*Nepenthes* pitchers of uniform age group (5-8 days after opening) were collected from the conservatories in the morning hours and their physical parameters were noted (Supplementary Table S1). Pitcher lids were carefully plucked from the peristome-lid intersections; peristomes were circularly (externally) cut from the pitchers using sharp blades. Whole peristomes and lids (without slicing) were rinsed (individually, separately) with water (5 ml, 15 min each, 5-8 times), washes were pooled and concentrated to dryness using a R-210 rotary evaporator (Buchi, Switzerland) at 37°C under reduced pressure. The completion of nectar extraction was ensured by checking the absence of sugars in water washes by thin layer chromatography (TLC). Peristome and lid EFN yields were noted. This isolation protocol was repeated to obtain adequate quantity of EFNs for the experiments.

Extrafloral nectar in the chosen *Nepenthes* species/hybrids are mostly adhered to the pitcher peristome and lid surfaces, and other methods of nectar sampling (Bauer *et al*., 2009; Bauer *et al*., 2016) are not feasible. Therefore, in this study, *Nepenthes* peristomes/lids were water-rinsed repeatedly, water in the washings was removed under vacuum and the ‘dry nectar’ was denoted as ‘extrafloral nectar’ (‘EFN’).

### Sugars

Quantitative estimation of glucose (Glc), fructose (Fru), and sucrose (Suc) contents in EFNs of *Nepenthes* pitcher peristomes and lids was carried out by HPTLC-densitometry (CAMAG, Switzerland), comprising of an automatic Linomat V sample applicator, twin trough plate development chamber, TLC Scanner 3, TLC Visualizer 2 and VisionCATS software 3.2, using pre-treated silica gel plates (60 F_254,_ 20 x 10 cm, 0.2 mm thickness; E. Merck, Germany). These TLC plates were modified by dipping in 0.2 M potassium dihydrogen orthophosphate solution (aqueous), wet plates were dried at 90°C for 10 min, and allowed to cool to room temperature in a desiccator (Govindan *et al*., 2016).

Stock solutions of Glc, Fru and Suc standards (Sigma Aldrich, USA; 10 mg/2.4 ml each) were prepared in 1:2 water-MeOH (v/v), and their different concentrations were applied onto the pre-treated silica gel TLC plates with the Linomat V sample applicator, fitted with a microsyringe, under N_2_ flow (application rate 100 nl/s, space between two bands 12.1 mm). Plates were developed three times, each up to 80 mm in a paper-lined twin trough plate development chamber, equilibrated with the mobile phase, acetonitrile:water (8.5:1.5, v/v; 20 ml). The mobile phase was replaced with fresh solvent for each run, and the plates were thoroughly dried in between the runs. After the third run the plates were dried, derivatized using anisaldehyde-sulphuric acid reagent, heated in a hot air oven at 110°C for 5 min and scanned densitometrically at 570 nm using TLC Scanner 3 (slit dimension 6.00 × 0.45 mm, scanning speed 20 mm/s) equipped with VisionCATS software 3.2. Calibration curves of standard sugars (Glc, Fru, Suc: 0.28, 0.69, 1.39, 2.09, 2.78, 3.48, 4.17 μg) were obtained by plotting their quantities *versus* areas (Glc: y = 0.0037x + 0.001, r^2^ = 0.99; Fru: y = 0.0027x + 0.0008, r^2^ = 0.99; Suc: y = 0.0039x + 0.0019, r^2^ = 0.99).

*Nepenthes* pitcher peristome and lid EFNs (1 mg each) were dissolved in 0.8:0.2 MeOH-water (v/v, 1 ml), 10 μl of these solutions were applied to modified silica gel TLC plates as 8 mm wide bands and the plates were developed, derivatized and scanned as in the case of the sugar standards; Glc, Fru and Suc contents in EFNs were quantified using the calibration curves. *Nepenthes khasiana* (n = 6), *N. mirabilis* (n = 2), *N. mirabilis* × *N. khasiana* (n = 5), *N. mirabilis* × *N. rafflesiana* (n = 3) peristome and lid EFNs were analyzed and their Glc-Fru-Suc contents (%) are listed in Supplementary Table S1.

### Amino acids

Amino acids in *Nepenthes* pitcher peristome and lid EFNs were analyzed by HPTLC-densitometry (CAMAG, Switzerland). Stock solutions (1 mg/ml) of twenty standard amino acids (Ala, Arg, Asn, Asp, Cys, Glu, Gln, Gly, His, Ile, Leu, Lys, Met, Phe, Pro, Ser, Thr, Trp, Tyr, Val) (Fluka Analytical, USA) were prepared in 0.9:0.1 MeOH-water (v/v, 1 ml) and applied (2 μl, each) onto silica gel TLC plates (60 F_254,_ 20 x 10 cm, 0.2 mm thickness; E. Merck, Germany) with the Linomat V sample applicator, fitted with a micro syringe, under N_2_ flow (application rate 100 nl/s, space between two bands 12.1 mm). Plates were developed up to 80 mm in the twin trough plate development chamber saturated with butanol:acetic acid:water (12:3:5, v/v, 20 ml). The developed plates were dried, derivatized using ninhydrin reagent, heated at 110°C for 5 min, and scanned densitometrically at 570 nm using TLC Scanner 3 (slit dimension 6.00 × 0.45 mm, scanning speed 20 mm/s) equipped with VisionCATS software 3.2. *Nepenthes* pitcher peristome and lid EFNs (1 mg each; *N. khasiana* n = 6, *N. mirabilis* n = 3, *N. mirabilis* × *N. khasiana* n = 2, *N. mirabilis* × *N. rafflesiana* n = 3) were dissolved in 0.8:0.2 MeOH-water (v/v, 1 ml), 5 and 10 μl of these solutions were applied to silica gel TLC plates as 8 mm wide bands, plates were developed, derivatized and scanned as in case of the amino acid standards (Supplementary Fig. S1). Amino acids in EFNs were also tested by ultra-fast liquid chromatography (UFLC) (Supplementary Fig. S2) and the ninhydrin test.

### Proteins

Total protein contents in *Nepenthes* peristome and lid EFNs (*N. khasiana* n = 6, *N. mirabilis* n = 3, *N. mirabilis* × *N. khasiana* n = 2, *N. mirabilis* × *N. rafflesiana* n = 3) were determined colorimetrically using the Bradford protein assay kit (Thermo Scientific, USA) with bovine serum albumin (BSA) standard (Bradford, 1976).

### Vitamin C

Vitamin C contents in *Nepenthes* species/hybrid peristome and lid EFNs (*N. khasiana* n = 6, *N. mirabilis* n = 3, *N. mirabilis* × *N. khasiana* n = 2, *N. mirabilis* × *N. rafflesiana* n = 3) were estimated titrimetrically by indophenol method with ascorbic acid as the standard (Nielson, 2019).

### Fatty acids

Peristome (145.0 mg) and lid (78.4 mg) EFNs isolated from *N. khasiana* pitchers were dissolved in 10 ml water (each), partitioned with hexane (10 ml × 5) separately and concentrated in a rotary evaporator (Buchi, Switzerland) at 30°C under reduced pressure to obtain their hexane (peristome 2.4 mg, lid 0.6 mg) fractions. These hexane fractions were dissolved (separately) in hexane (1 ml), added with methanolic NaOH (2 N, 200 μl) and heated at 50°C (20 s) followed by shaking (10 s). Methanolic HCl (2 N, 200 μl) was added to these mixtures, upper layers were collected and dried over anhydrous sodium sulphate (de Melo Soares *et al*., 2022). These esterified fractions (1 μl each) diluted in diethyl ether (1:10 dilution, separately) were injected onto a QP2020C NX Gas Chromatograph Mass Spectrometer (Shimadzu, Japan), fitted with an SH-Stabilwax (Crossbond Carbowax polyethylene glycol) capillary column (30 m x 0.32 mm i.d., film thickness 0.25 µm), coupled with a single quadrupole 8030 series mass selective detector. GC-MS parameters: injector temperature 240°C, oven temperature 60-250°C (3°C/min), ion source temperature 240°C, and interphase temperature 260°C. Individual compounds were identified using NIST 17 and Wiley 275 library search. Palmitic acid, stearic acid and *cis*-11-octadecenoic acid standards and their methyl esters were analyzed in GC-MS under similar conditions, and the results are listed as relative abundance (%). Repeated fractionation, FAME-derivatization and GC-MS analysis (n = 3) showed similar fatty acid patterns in *N. khasiana* EFNs.

### Minerals

Mineral (Ag, Al, B, Ba, Bi, Ca, Cd, Co, Cr, Cu, Fe, Ga, In, K, Mg, Mn, Na, Ni, Pb, Tl, Zn) contents in *N. khasiana* peristome and lid EFNs were analyzed by Avio 200 Inductively Coupled Plasma Optical Emission Spectrometer (PerkinElmer, USA) at a spectral range 165-900 nm with a resolution of < 0.009 nm at 200 nm. Fourth-generation 40 MHz, free-running solid-state RF generator, which is adjustable from 1000 to 1500 watts in 1-watt increments was used. ICP multi-element standard IV was used as the standard, and a five-level calibration was carried out. *N. khasiana* peristome and lid EFNs (0.1 g each, separately) were dissolved in 15 ml ultrapure water, filtered, drops of nitric acid added, and analyzed in triplicate (Supplementary Table S2).

### Volatiles

Headspace GC-MS analyses of fresh *N. khasiana* specimens *viz.*, tendril (0.54-0.59 g), peristome (0.99-1.38 g), lid (0.91-1.28 g), slice of the mid interior portion of peristome (nectary, 0.39-0.67 g), pitcher fluid (50 μl), and plumbagin standard (1 mg) were carried out on a QP2020C NX Gas Chromatograph Mass Spectrometer (Shimadzu, Japan) attached to an HS20 Headspace Sampler (Shimadzu, Japan), fitted with an SH-Rxi-5Sil MS (Crossbond similar to 5% diphenyl/95% dimethyl polysiloxane) capillary column (30 m, 0.25 mm i.d., film thickness 0.25 µm). A droplet of EFN collected from the inner side *N. khasiana* lid using a syringe needle in early morning hours was also analyzed. Vial containing each sample was heated at 65°C for 10 min at shaking level 3 and pressurized up to 60 kPa. GC oven temperature: 50-200°C (10°C/min); ionization mode: electron impact ionization (EI) 70 eV; data processing: GC-MS solution Ver. 4. Individual compounds were identified using NIST 17 and Wiley 275 library search, and the results are presented as relative abundance (%). *Nepenthes khasiana* tendril, peristome, lid, nectaries and pitcher fluid on repeated headspace GC-MS analyses (n = 3) gave identical results.

### CHN analysis

Carbon, hydrogen and nitrogen contents of *N. khasiana* peristome and lid EFNs (n = 3 each) were analyzed on a 2400 Series II CHNS Organic Elemental Analyzer (PerkinElmer, USA). Freeze-dried EFNs (1-2 mg) were used for the analysis, chromatographic responses were calibrated against pre-analyzed cystine standards, and CHN contents are reported in weight percent.

### Prey spectrum

Prey composition of *N. khasiana* pitchers was studied by collecting the pitcher contents from 2 to 14 days after opening (DAO). Destructive sampling was done at regular intervals of 2, 4, 6, 8, 10, 12 and 14 DAO of the pitchers. Six pitchers each (total 42) were sampled at these time intervals from *N. khasiana* plants at JNTBGRI conservatory. The contents of the pitchers were washed in double distilled water, sorted out using a stereo binocular microscope (Leica MZ6, Germany) and preserved in 75% alcohol for further identification. The insects were identified up to order level and data are listed in Supplementary Table S3 (Fig. 1).

**Fig. 1.**
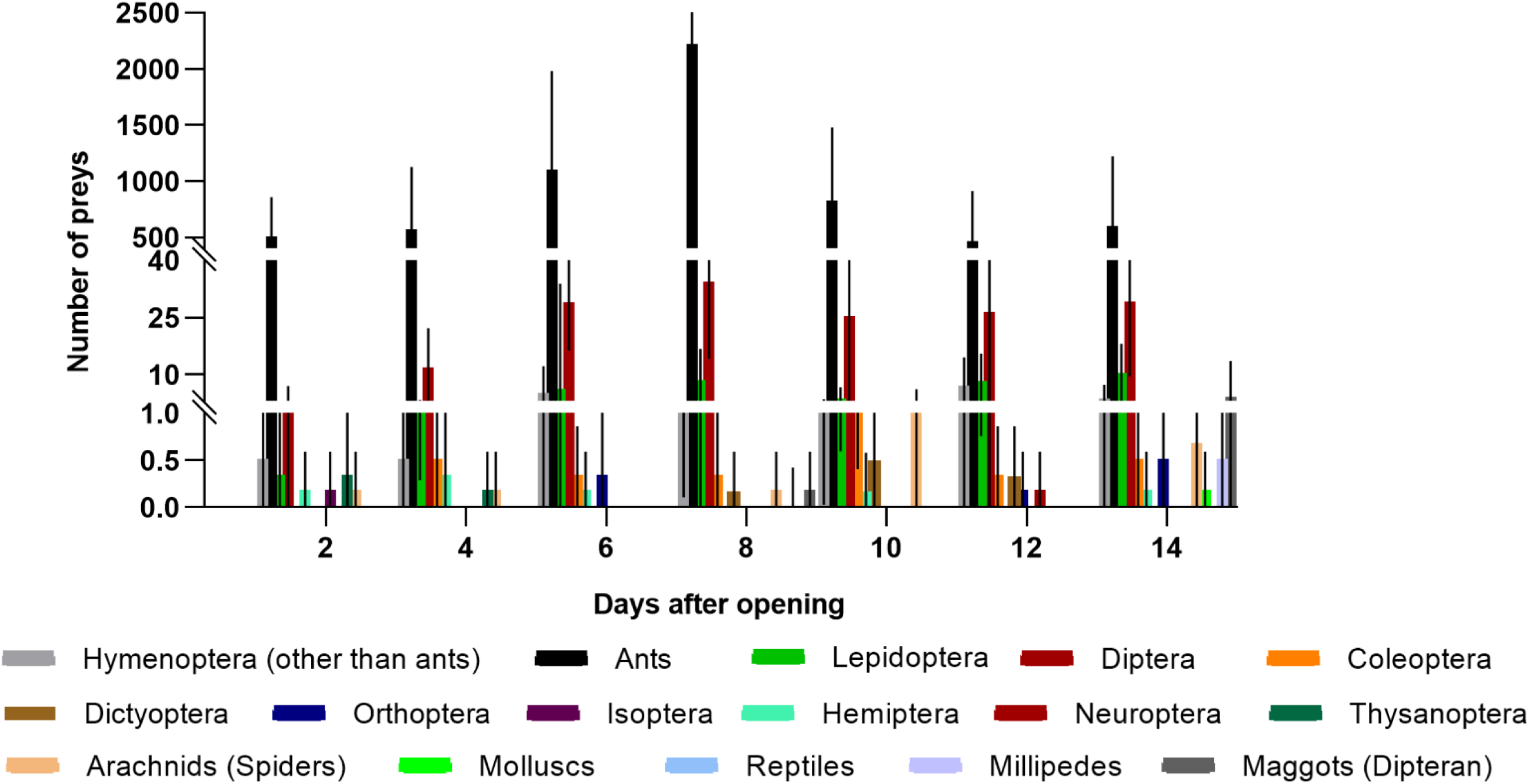
Prey capture by *Nepenthes khasiana* pitchers. Insects and other organisms trapped in *N. khasiana* pitchers from 2 days to 14 days after opening (data presented as average prey capture per pitcher, n = 6; details in Table S3).

### Ants

Two commonly visiting ant species, *viz.*, *Anoplolepis gracilipes* Smith (yellow crazy ant, long-legged ant), *Monomorium minutum* Mayr. (little black ant), were collected from opened *N. khasiana* pitchers. For this, *N. khasiana* pitcher fluids with prey capture were transferred to petri dishes, fluid volume(s) were reduced using micropipettes, and these two species of test (EFN-fed) ants were picked with the help of a handheld lens and light source. The same two species of control ants were captured from the stem and leaf surfaces of *N. khasiana*. Test (with no pitcher fluid) and control ants were carefully transferred to suitable containers and kept at 4°C for the experiments.

### AChE inhibition by EFN-fed ants (*in vivo*)

Acetylcholine iodide (AChI), bovine erythrocyte cholinesterase (AChE), 5,5′-dithiobis(2-nitrobenzoic acid) (DTNB) and quinidine sulphate were procured from Sigma-Aldrich (MO, USA). Test ants (*A. gracilipes*, *M. minutum*; 20 each) collected from the pitcher fluids of *N. khasiana* were transferred to petri dishes (separately). Crude homogenates of ants were prepared in Eppendorf tubes using a manual homogenizer at 4°C in phosphate buffer (1 ml, 50 mM, pH 7.0). Homogenates were centrifuged at 12,000 × g for 15 min at 4°C and supernatants were separated; these supernatants of both test ants were used to prepare the AChE enzyme sources at a concentration of 400 units/l as described by Chinyere and Christos (2019). Similarly, supernatants of control ants were also prepared. *Nepenthes khasiana* pitcher fluid with prey capture was filtered to remove the ants and other captured preys, and this light yellow fluid at different volumes was directly tested for AChE inhibition (Supplementary Table S4).

For both test and control ant homogenates, reaction mixes were prepared by combining the substrate (AChI, 100 μl, 136 µg/ml), enzyme source (supernatant, 50 µl, 5 units/l) and DTNB (60 μl, 0.4 mg/ml), made up to 500 μl using phosphate buffer (pH 8.0). For pitcher fluid, reaction mixes were prepared by combining the substrate (AChI, 100 μl, 136 µg/ml), enzyme (AChE, 50 µl, 5 units/l) and DTNB (60 μl, 0.4 mg/ml). Pitcher fluid at different volumes (1 to 100 µl) was added to the respective wells, and the final volumes were made up to 500 μl using phosphate buffer (pH 8.0). The samples were incubated at 30°C for 15 min, and quinidine sulphate (0.1%) was added to stop the reactions. Absorbances were measured at 412 nm at 0 min and 10 min intervals for 4 h using an iMark^TM^ Microplate Absorbance Reader (Bio-Rad, Japan) (Ellman *et al*., 1961; Gerczei and Pattison, 2015). Donepezil hydrochloride (2.5-1000 µg/ml) served as the drug control (Colović *et al*., 2013). All samples were assayed in triplicate (Fig. 2, Supplementary Table S4).

**Fig. 2.**
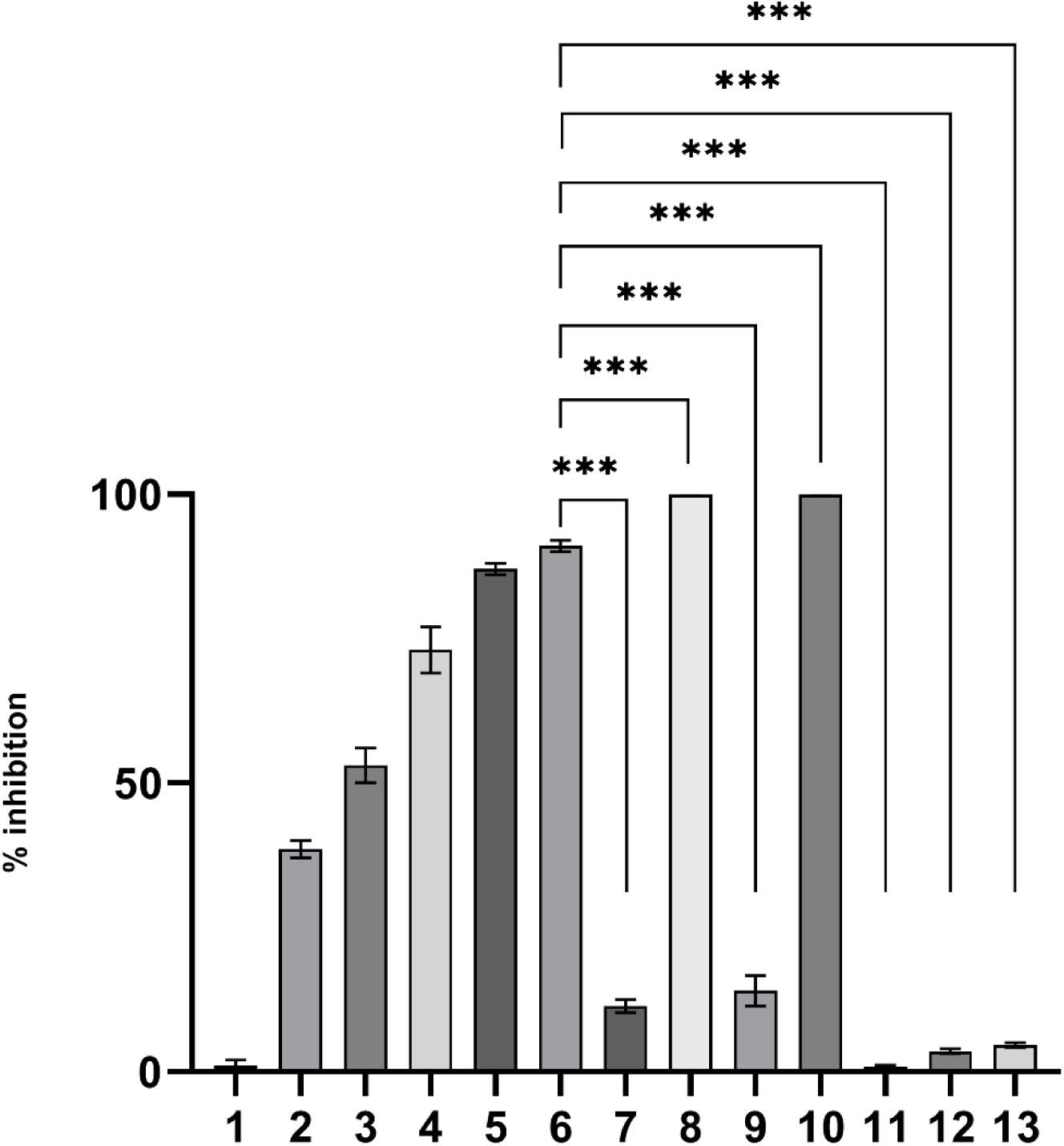
AChE inhibition by homogenates of captured ants, *M. minutum* and *A. gracilipes, (in vivo)* and pitcher fluid col­lected from *N. khasiana* pitchers. **1:** substrate (AChl) + AChE; **2:** AChl + donepezil (2.5 pg/ml) + AChE; **3:** AChl + donepezil (5.0 pg/ml) + AChE; 4: AChl + donepezil (10 pg/ml) + AChE; 5: AChl + donepezil (100 pg/ml) + AChE; **6:** AChl + donepezil (1000 pg/ml) + AChE; 7: AChl + control-enzyme source (non-EFN fed ant *M. minutum* homogenate); 8: AChl + test-enzyme source (EFN-fed ant *M. minutum* homogenate); 9: AChl + control-en­zyme source (non-EFN fed ant *A. gracilipes* homogenate); **10:** AChl + test-enzyme source (EFN-fed ant *A. gracilipes* homoge­nate); **11:** AChl + pitcher fluid (1 pl) + AChE; **12:** AChl + pitcher fluid (5.0 pl) + AChE; **13:** AChl + pitcher fluid (10 pl) + AChE; AChl (136 pg/ml) and donepezil are substrate and control, respectively; AChE, control and test ant homogenates, 5 units/l. *p* values were calculated using one-way ANOVA and Dunnett’s multiple comparisons test using GraphPad Prism 8.0.2; values are mean ± S.D; n = 3; *\*\*\*p* < 0.0001 (compared to 6); data listed in Table S4

A standard curve was generated by preparing reaction mixes consisting of varying concentrations of the substrate (AChI, 136 µg/ml), enzyme (AChE, 50 µl, 5 units/l) and DTNB (60 μl, 0.4 mg/ml), made up to a final volume of 500 µl with phosphate buffer (pH 8.0) and subjected to similar measurements. Protein contents of enzyme sources were determined by the Lowry assay (Lowry *et al*., 1951), and percentage AChE inhibitions were calculated using the standard curve (Ellman *et al*., 1961).

### Activity guided isolation

AChE inhibiting entity in *N. khasiana* EFNs was isolated by activity guided isolation. Peristome (270.4 mg) and lid (249.1 mg) EFNs isolated from 48 pitchers were subjected to column chromatography separately over silica gel, and the columns were eluted with hexane, hexane:CHCl_3_, CHCl_3_:MeOH and MeOH. Similar fractions were pooled together based on TLC profiles, which resulted in six major fractions from each column (peristome EFN fractions: NPF1 to NPF6; lid EFN fractions: NLF1 to NLF6). These major fractions were tested for AChE inhibition, and NPF5 and NLF4 showed significant activity. TLC profiles of these two active fractions displayed a major band at the same retention factor. To isolate this compound, peristome EFN (458.0 mg, isolated from 59 pitchers) was subjected to silica gel column chromatography, eluted with hexane, hexane:CHCl_3_, CHCl_3_, CHCl_3_:MeOH and MeOH. Six major fractions were obtained, of which fifth fraction yielded the active component (3.1 mg only).

Higher amounts of the active principle was isolated from the peristomes of *N. khasiana* pitchers. Peristomes of 30 pitchers (18.6 g) were extracted with distilled water (100 ml), and the filtered extract was evaporated to dryness using a rotary evaporator at 37°C. This extract (1.2 g) was subjected to silica gel column chromatography. Similar fractions were pooled together, and the sixth fraction yielded the impure active compound (10.0 mg). These extraction-isolation steps were repeated three times to obtain 23.5 mg of the impure fraction. Re-chromatography of this fraction over silica gel by eluting with hexane:CHCl_3_ yielded the active compound (14.0 mg) (details of the isolation protocol are given in the Supplementary Information).

### Spectroscopy of active principle

1H-, 13C-NMR (400 MHz, Bruker, Germany), LC-HRMS (Thermo Fisher Scientific, USA), FTIR (Thermo Scientific, USA) spectra and optical rotation (P2000 Polarimeter, Jasco, Japan) of the active molecule were recorded and its structure was elucidated (Hanson *et al*., 1981) (Supplementary Table S5, Supplementary Figs. S3-S5).

### LC-MS metabolic profiling

Stock solutions (1 µg/µl) of *N. khasiana* peristome and lid EFNs were prepared in water and filtered through 0.45 µm syringe filters. Metabolic profiling of the EFNs was carried out using a LC-2050C 3D i-Series Liquid Chromatograph-Mass Spectrometer (Shimadzu, Japan) equipped with a C-18 HPLC Shim-pack (5 µm, 250 x 4.6 mm) column. Peristome and lid EFNs (25 µl each, 1 µg/µl) were separately injected onto the LC-MS system and a gradient mobile phase with water (solvent A, HPLC grade) and acetonitrile (solvent B, HPLC grade) both containing 0.1% formic acid (flow rate 0.3 ml/min, column temperature 40°C) was applied (initial elution 0% B, increased to 5.05% B in 8.80 min, 17.68% B in 30.70 min, 30.30% B in 52.60 min, 37.88% B in 65.80 min, 38.13% B in 66.20 min and finally to 50.51% B in 87.7 min). Data were acquired on an LCMS-2050 Mass Spectrometer (Shimadzu, Japan); metabolites were detected in positive ionization mode (MS parameters: scan range 100-400 m/z, gas temperature 450°C, drying gas 5 l/min, nebulizer gas flow 2 l/min) (Supplementary Fig. S6).

### AChE inhibition by EFN, active principle (*in vitro*)

Extrafloral nectars from *N. khasiana*, *N. mirabilis* and hybrid (*N. mirabilis × N. khasiana*, *N. mirabilis × N. rafflesiana*) peristomes and lids, active fractions of *N. khasiana* EFNs and active compound, (+)-isoshinanolone, were tested for AChE inhibition activity. Briefly, a mix for each reaction was prepared by combining the substrate (AChI, 100 μl, 136 µg/ml), enzyme (AChE, 50 µl, 5 units/l) and DTNB (60 μl, 0.4 mg/ml) in a 12 well plate. Thereafter, EFN or active fraction or compound at various concentrations were added to the respective wells, and the final volumes were made up to 500 μl using phosphate buffer (pH 8.0). The samples were incubated at 30°C for 15 min, and quinidine sulphate (0.1%) was added to stop the reactions. Absorbances were measured at 412 nm at 0 and 10 min intervals for 4 h. Appropriate controls like (substrate (AChI) + enzyme source), (substrate (AChI) + AChE), (drug (donepezil) + substrate (AChI) + enzyme source) and (drug (donepezil) + substrate (AChI)+ AChE) were also tested; all samples were assayed in triplicate (Fig. 3, Supplementary Tables S6, S7; Supplementary Figs. S7, S8). Similarly, plumbagin isolated from *N. khasiana* (whole plant; Raj *et al*., 2011) was tested for its AChE inhibition (Supplementary Table S8, Supplementary Fig. S9).

**Fig. 3.**
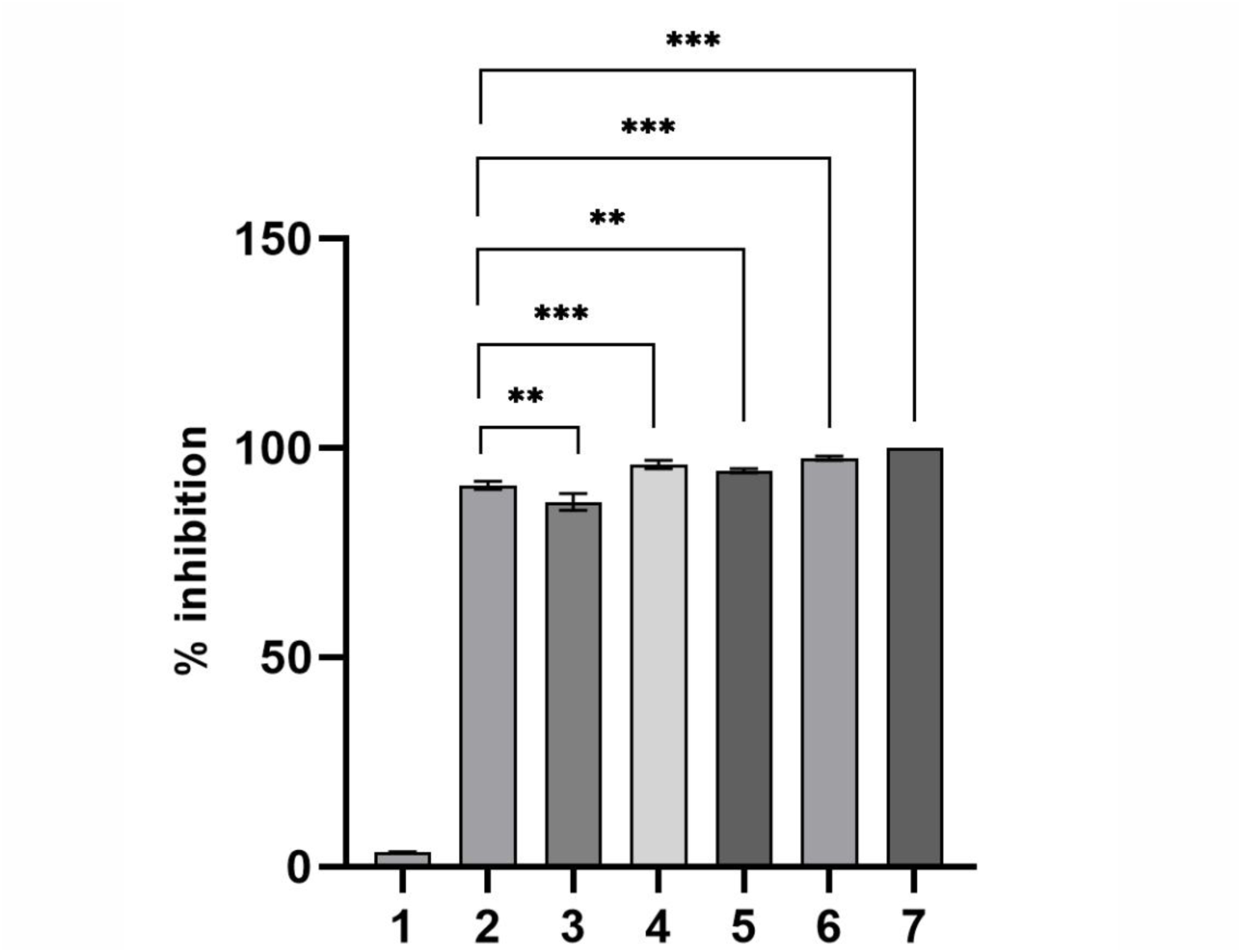
AChE inhibition of *N. khasiana* EFNs and the active principle, (+)-isoshinanolone *(in vitro)*. **1:** AChl + AChE; 2: AChl + donepezil (1000 pg/ml) + AChE; 3: AChl + peristome EFN (1000 pg/ml) + AChE; 4: AChl + lid EFN(1000 pg/ml) + AChE; 5: AChl + (+)-isoshinanolone (10 pg/ml) + AChE; 6: AChl + (+)-isoshinanolone (100 pg/ml) + AChE; 7: AChl + (+)-isoshinanolone (1000 pg/ml) + AChE; AChl (136 pg/ml) and donepezil are substrate and control, respectively; AChE, 5 units/l. *p* values were calculated using one-way ANOVA and Sidak’s multiple comparisons test using GraphPad Prism 8.0.2; values are mean ± S.D; n = 3; **p = 0.0010, 0.0034; *\*\*\*p* < 0.0001 (compared to 2); data listed in Table S6.

### AChE inhibition by EFN-, (+)-isoshinanolone-fed ants (*in vivo*)

*Anoplolepis gracilipes* ants were collected from their natural habitats at the campus sites of JNTBGRI, and 20 ants each were transferred to plastic jars with ventilation holes on their lids. These ants were allowed to feed 1 ml each of sugar mix (Glc 0.60 mg + Fru 1.03 mg + Suc 0.55 mg, in distilled water), (sugar mix + donepezil), (sugar mix + (+)-isoshinanolone), *N. khasiana* peristome and lid EFNs at various concentrations as listed in Supplementary Table S9. These experimental jars were kept in a temperature (22-24°C)-controlled room overnight. Suc:Glc:Fru ratio in the sugar mix was the same as in *N. khasiana* peristome EFN (Supplementary Table S1). Crude homogenates of these test and control (unfed) ants were prepared separately, 12 h after feeding, in Eppendorf tubes using a manual homogenizer at 4°C in phosphate buffer (1 ml, 50 mM, pH 7.0), centrifuged at 12,000 × g for 15 min at 4°C and supernatants were separated. These supernatants were used to prepare the AChE sources at a concentration of 400 units/l, and AChE inhibition assays were carried out as described earlier (Supplementary Table S9, Fig. 4).

**Fig. 4.**
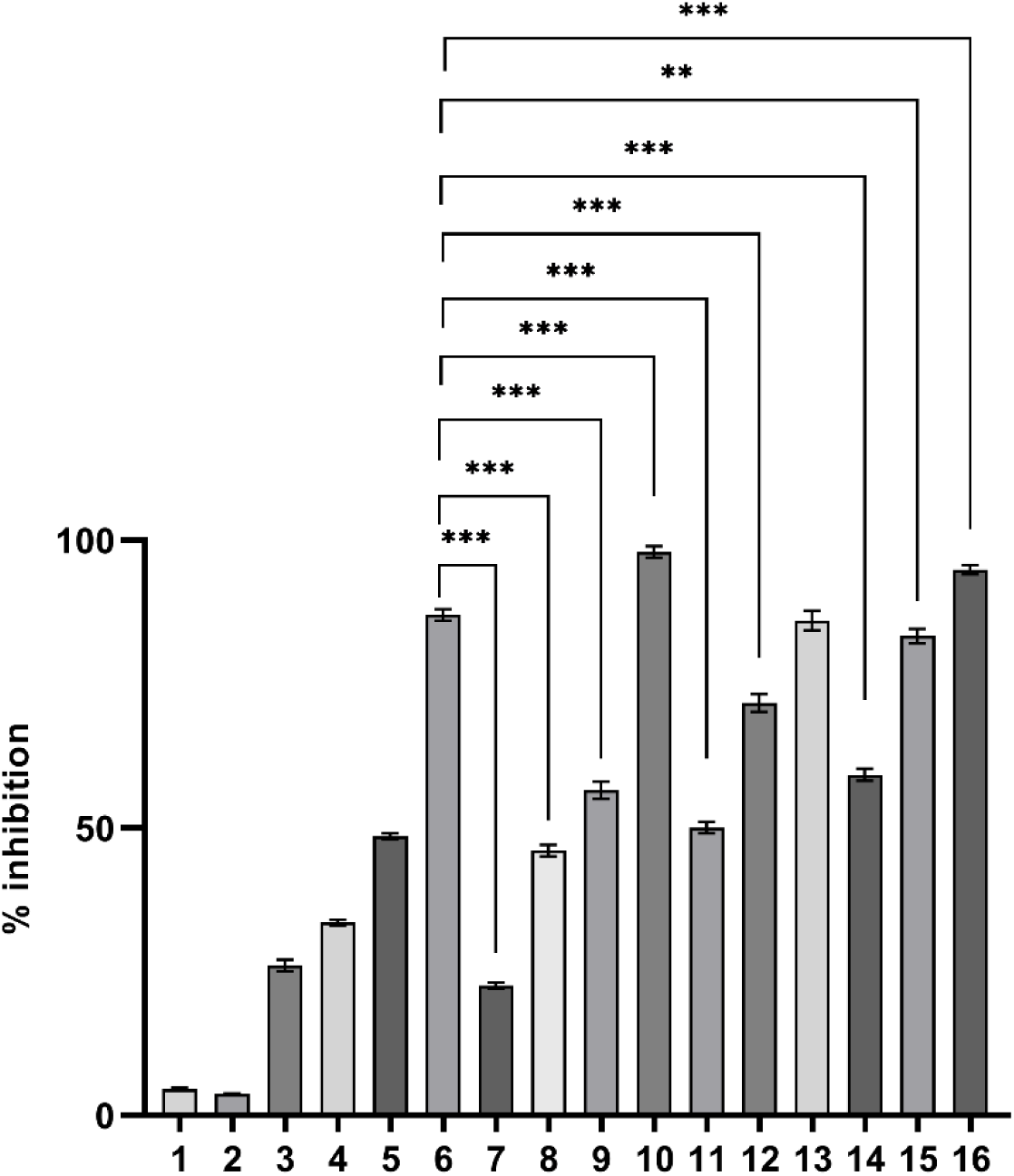
AChE inhibition by *N. khasiana* EFN– and (+)-isoshi-nanolone-fed ant, A. *gracilipes (in vivo).* 1: AChl + unfed ant homogenate; 2: AChl + sugar mix (Sue 0.55 mg + Glc 0.60 mg + Fru 1.03 mg)-fed control ant homogenate; 3: AChl + (sugar mix + donepezil 1.25 pg/ml)-fed ant homogenate; 4: AChl + (sugar mix + donepezil 2.5 pg/ml)-fed ant homogenate; **5:** AChl + (sugar mix + donepezil 5.0 pg/ml)-fed ant homogenate; **6:** AChl + (sugar mix + donepezil 10 pg/ml)-fed ant homogenate; 7: AChl + (sugar mix + (+)-isoshinanolone 1.25 pg/ml)-fed ant homoge­nate; 8: AChl + (sugar mix + (+)-isoshinanolone 2.5 pg/ml)-fed ant homogenate; 9: AChl + (sugar mix + (+)-isoshinanolone 5 pg/ml)-fed ant homogenate; 10: AChl + (sugar mix + (+)-isoshi-nanolone 10 pg/ml)-fed ant homogenate; 11: AChl + peristome EFN (10 pg/ml)-fed ant homogenate; 12: AChl + peristome EFN (100 pg/ml)-fed ant homogenate; **13:** AChl + peristome EFN (1000 pg/ml)-fed ant homogenate; **14:** AChl + lid EFN (10 pg/m-l)-fed ant homogenate; 15: AChl + lid EFN (100 pg/ml)-fed ant homogenate; 16: AChl + lid EFN (1000 pg/ml)-fed ant homoge­nate. AChl (136 pg/ml) and donepezil are substrate and control, respectively; control and test ant homogenates, 5 units/l each, ***p*** values were calculated using one-way ANOVA and Dunnett’s multiple comparisons test using GraphPad Prism 8.0.2; values are mean ± S.D; n = 3; ***\*\*p*** = 0.0014; ***\*\*\*p*** < 0.0001 (compared to 6); data listed in Table S9.

In similar biotests (24 h, *in vivo*), *A. gracilipes* ants (20 each) were fed with sugar mix (only), *N. khasiana* peristome EFN (1000 µg/ml), sugar mix + (+)-isoshinanolone (10 µg/ml) and sugar mix + donepezil (10 µg/ml) (1 ml each), and their behavioral changes were carefully observed every hour for 24 h. After 24 h feeding, ants in all four groups and control (water only-fed) ants were harvested, and their AChE inhibition assays were performed in triplicate.

## Statistical analysis

In AChE inhibition assays, quantitative data are presented as the mean ± S.D. of three replicates. Statistical analysis was carried out by one-way ANOVA and Sidak’s/Dunnett’s multiple comparison tests using GraphPad Prism 8.0.2 software.

## Results

### EFN chemistry

*Nepenthes khasiana* pitcher peristomes and lids yielded 10.00 ± 8.48 mg (n = 6) and 6.23 ± 4.94 mg (n = 6) EFNs, respectively. EFNs showed Glc-Fru-Suc patterns (peristome, Glc: 6.03 ± 3.38%, Fru: 10.27 ± 6.85%, Suc: 5.53 ± 3.38%, n = 6; lid, Glc: 7.76 ± 5.40%, Fru: 15.80 ± 7.00, Suc: 3.67 ± 3.47%, n = 6), along with few other minor sugars (Supplementary Table S1). EFNs of *N. mirabilis* and the two hybrids, *N. mirabilis × N. khasiana*, *N. mirabilis × N. rafflesiana*, also displayed sugar patterns (Supplementary Table S1). *Nepenthes mirabilis*, with smaller pitchers, showed relatively low nectar yields (peristome: 1.85 ± 0.35 mg (n = 2), lid: 1.95 ± 0.21 mg (n = 2)) and sugar patterns as (peristome, Suc: 1.15 ± 1.62% (n = 2); lid, Suc: 3.70 ± 5.23% (n = 2)). Glc and Fru were absent in *N. mirabilis* EFNs. Similarly, either one or all three sugars were absent in a few EFNs (Supplementary Table S1).

Only very low levels of proteins were found in *N. khasiana* (peristome: 1.85 ± 2.25%, w/w; lid: 1.20 ± 2.86%, w/w, n = 6) and *N. mirabilis × N. khasiana* (peristome: 0.61 ± 0.05% w/w; lid: 0.22 ± 0.02%, w/w, n = 2) EFNs. No free proteins were detected in *N. mirabilis* and *N. mirabilis × N. rafflesiana* peristome, lid EFNs. *Nepenthes khasiana* peristome and lid EFNs were analyzed for free amino acids using HPTLC-densitometry, and none were detected (Supplementary Fig. S1); ninhydrin tests also gave similar results. In UFLC analysis, only trace of methionine (Met; 0.0008%) was detected in the peristome EFN of *N. khasiana*, and none of the amino acids were detected in its lid EFN (Supplementary Fig. S2). That is, free amino acids are minimal or below detection levels in *N. khasiana* EFNs. On similar assays, *N. mirabilis* and the two test hybrid EFNs also showed below detection limits of amino acids. Vitamin C was also minimal (or absent) in the EFNs of *N. khasiana*, *N. mirabilis* and the two hybrids. *Nepenthes khasiana* EFNs on FAME-derivatization and gas chromatographic analysis showed C16:0, C18:1, C18:0 fatty acids (peristome: palmitic acid methyl ester (C16:0 derivative) 43.34%, *cis*-11-octadecenoic acid (C18:1) 23.33%, palmitic acid (C16:0) 11.96%; lid: *cis*-11-octadecenoic acid (C18:1) 55.79%, palmitic acid (C16:0) 19.73%, stearic acid (C18:0) 13.25%). Mineral analysis revealed *N. khasiana* peristome and lid EFNs as rich in sodium (peristome, lid: 24.21 ± 4.84, 68.67 ± 13.74 ppm), potassium (48.14 ± 8.97, 159.8 ± 31.96 ppm), calcium (7.49 ± 1.18, 23.42 ± 3.40 ppm) and magnesium (3.83 ± 0.77, 12.98 ± 2.59 ppm); toxic minerals such as cadmium, chromium and lead were below detection levels (Supplementary Table S2). *Nepenthes khasiana* EFNs displayed mild green fluorescence at UV 366 nm. Again, its EFNs (peristome: C 34.65 ± 1.71%, N 0.43 ± 0.06%, n = 3; lid: C 33.90 ± 0.73%, N 0.43 ± 0.01%, n = 3) showed high C:N ratios.

### AChE inhibition, toxic principle

Two nectar-fed ant species (their homogenates, 5 units/l each, *in vivo*), *A. gracilipes* and *M. minutum*, collected from *N. khasiana* pitcher fluids displayed near 100% AChE inhibition (99.9%), better than donepezil (1000 µg/ml, 91.0%), whereas the homogenates of control ants captured from *N. khasiana* stem, leaf surfaces showed only minimal enzyme inhibition (14.0-15.3%) (Fig. 2, Supplementary Table S4). *N. khasiana* pitcher fluids (with prey capture) when directly tested displayed 0.9% (1 µl), 1.4% (2.5 µl), 3.6% (5.0 µl), 4.6% (10.0 µl), 19.7% (25.0 µl), 59.3% (50.0 µl) and 94.7% (100.0 µl) AChE inhibition (Fig. 2, Supplementary Table S4). The maximum probable drinking volumes of pitcher fluid for the test ants (*A. gracilipes*, 1-10 µl) did not show any significant AChE inhibition (Fig. 2), but higher volumes (50.0-100.0 µl) showed better activity (Supplementary Table S4).

*Nepenthes khasiana* peristome and lid EFNs (1000 µg/ml) displayed 87.0% and 96.0% AChE inhibition (*in vitro*), respectively (Supplementary Table S7); and their chromatographic fractions showed varying levels of (few fractions ˃ 80%) inhibition. The active constituent from the most active peristome and lid EFN fractions was isolated by column chromatography and characterized by spectroscopy as the naphthoquinone derivative, (+)-isoshinanolone (Supplementary Table S5; Supplementary Figs. S3-S5). Phytochemical analysis of *N. khasiana* peristome tissues also yielded (+)-isoshinanolone. Moreover, LC-MS-metabolic profiling detected (+)-isoshinanolone in *N. khasiana* EFNs (Supplementary Fig. S6). This active principle displayed 94.5%, 97.5% and 100.0% AChE inhibition at 10, 100, 1000 µg/ml, comparable or above donepezil (Fig. 3, Supplementary Table S6). *Nepenthes mirabilis*, *N. mirabilis × N. khasiana* and *N. mirabilis × N. rafflesiana* peristome and lid EFNs at 10-1000 µg/ml showed 55.3-77.9%, 32.9-52.3% and 34.0-53.0% inhibitions, respectively (Supplementary Table S7, Supplementary Figs. S7, S8). (+)-Isoshinanolone was also detected in gas chromatographic profiles of peristome and lid EFNs of *N. khasiana*, *N. mirabilis* and the two study hybrids (Supplementary Fig. S10).

In direct biotests (12 h, *in vivo*), *N. khasiana* peristome EFN (10-1000 µg/ml)-, *N. khasiana* lid EFN (10-1000 µg/ml)-, sugar mix + (+)-isoshinanolone (10 µg/ml)– and sugar mix + donepezil (10 µg/ml)-fed *A. gracilipes* ants showed 52.5-87.0%, 62.5-96.0%, 98.0% and 87.0% AChE inhibition, respectively, whereas sugar-fed and untreated control ants did not show enzyme inhibition (Fig. 4, Supplementary Table S9); IC_50_s of (+)-isoshinanolone and donepezil were 4.45 and 5.25 µg/ml, respectively. Symptoms of cholinergic toxicity and even deaths were observed in EFN-, (+)-isoshinanolone– and donepezil-fed ants. In similar biotests (24 h, *in vivo*), major symptoms observed in *N. khasiana* peristome EFN (1000 µg/ml)-, sugar mix + (+)-isoshinanolone (10 µg/ml)– and sugar mix + donepezil (10 µg/ml)-fed *A. gracilipes* ants were recorded as slow movements or cramps (muscular weakness), enhanced grooming (head, body) activity, fallen upside down, spasms and even deaths (in (+)-isoshinanolone-fed ants, 10%) (Supplementary Videos S1, S2); similar behavioral patterns were observed in *A. gracilipes* ants on *N. khasiana* peristomes (Supplementary Video S5). Again, ants in all three treatment groups after 24 h displayed strong AChE inhibition (*N. khasiana* peristome EFN (1000 µg/ml) 95.7 ± 1.5%, sugar mix + (+)-isoshinanolone (10 µg/ml) 97.7 ± 2.5%, sugar mix + donepezil (10 µg/ml) 91.7 ± 5.0%), whereas sugar mix-fed (10.7 ± 1.2%; Supplementary Video S3) and control (water only-fed; Supplementary Video S3) (6.3 ± 0.5%) *A. gracilipes* ants showed only negligible activity.

### Volatiles

Volatiles play crucial role in prey attraction and herbivory. Our headspace-GC-MS analyses detected volatiles (in relative abundance %) in the prey capture regions of *N. khasiana*. Plumbagin was the major volatile constituent in *N. khasiana* tendril (72.55%), peristome (85.47%), lid (74.58%) and slices of the mid interior portion of peristome (nectary, 68.79%). A droplet of EFN collected from *N. khasiana* lid also showed traces (2.56%) of plumbagin; but it was not detected in its pitcher fluid. Plumbagin, the major volatile constituent in prey capture regions, in the whole plant (Raj *et al*., 2011), and in EFN (only traces) of *N. khasiana* also showed strong AChE inhibition (1.25-1000 µg/ml: 49.0-98.3%; IC_50_ 2.43 µg/ml) (Supplementary Table S8, Supplementary Fig. S9).

### Preys, herbivores

*Nepenthes khasiana* pitchers trap a wide spectrum of preys which include arthropods, molluscs and reptiles (Fig. 1, Supplementary Table S3). Arthropods found in the pitchers include Insecta, Arachnida and Diplopoda. Insects belonging to ten orders, *viz.*, Hymenoptera, Diptera, Lepidoptera, Coleoptera, Hemiptera, Dictyoptera, Orthoptera, Isoptera, Neuroptera and Thysanoptera were collected and identified. Ants (Formicidae, Hymenoptera) were the predominant preys (96.4% of total capture) in *N. khasiana* pitchers, followed by flies (Diptera, 2.4%). Insects belonging to Hymenoptera (bees, wasps) (0.3%), Lepidoptera (0.6%), Coleoptera (0.05%) and arachnids (spiders, 0.06%) were minimal in pitcher catches from 2 to 14 days after opening. Hemiptera, Orthoptera, Dictyoptera, Isoptera, Neuroptera, and Thysanoptera were noticed occasionally. It was observed that the prey abundance increased from two days after opening and reached maximum on the eighth day (Fig. 1, Supplementary Table S3). Herbivore attacks were rarely observed in *N. khasiana*.

## Discussion

### EFN chemistry

In angiosperms, extrafloral nectar primarily functions as a defense against herbivores; its chemical constituents are mainly carbohydrates along with amino acids, proteins, secondary metabolites, fatty acids and vitamins as minor entities (Nicolson, 2007, 2022; Heil, 2015). It is also a persistent food source for tiny arthropods. In *Nepenthes*, EFN secreted by their pitchers has been regarded as the major reward for ants and other arthropods that are lured to these traps (Bauer *et al*., 2008; Bennett and Ellison, 2009; Greenwood *et al*., 2011). Sugars in floral nectar (FN) meet the energy requirements of pollinators (Nicolson, 2022); similarly, sugars in *Nepenthes* EFNs act as an energy source to visiting arthropods. Unlike Suc being the major carbohydrate in most FNs (Nepi *et al*., 2018; Nicolson, 2022), *N. khasiana* peristome and lid EFNs are hexose-rich with Suc/(Glc+Fru) ratios of 0.34 and 0.16, respectively. *N. mirabilis × N. khasiana* EFNs are sucrose-rich with Suc/(Glc+Fru) ratios: peristome 1.65, lid: 1.60. EFNs of other *Nepenthes* species/hybrids showed inconsistent Glc-Fru-Suc patterns (even one or more of these sugars were absent). Variations in yields and Glc-Fru-Suc ratios observed between EFNs secreted by *Nepenthes* species and hybrid pitchers are due to varying enzyme activities (Lin *et al*., 2014; Heil, 2015; Minami *et al*., 2021; Nicolson, 2022). Fru, the major sugar in *N. khasiana* EFNs, is the sweetest of naturally occurring carbohydrates, and it has the highest solubility of most sugars. All three sugars (Glc, Fru, Suc) are hygroscopic, Fru being the most; they tend to adsorb and retain moisture from the environment (Hanover and White, 1993), which is one of the factors causing peristome surface wetness leading to high prey capture rates (Bauer *et al*., 2008; Sujatha *et al*., 2023).

Strikingly, free amino acids were minimal (or < detection limit) and only very little protein was found in *Nepenthes* EFNs. Amino acids provide taste and nutritional benefits, while proteins function as preservatives in FNs (Grasso *et al*., 2015; Nicolson, 2022). Amino acids at trace levels undermine the nutritional gains of *Nepenthes* EFNs. Fatty acids (C16:0, C18:0, C18:1) and their derivatives were detected in *Nepenthes* EFNs; and they are used in resting metabolism in insects (Nicolson, 2022). A recent study showed that high CO_2_ enhanced fatty acid biosynthesis in the C3 species, *Arabidopsis thaliana* (*cac2-1* mutant) (Xi *et al*., 2023). Again, vitamin C, a vital nutrient which reduces oxidative damage (Nicolson, 2022), is minimal in *Nepenthes* EFNs. Khan and co-workers (2013) demonstrated the substantial decrease of vitamin C in tomato (*Lycopersicon esculentum*, C3 species) under elevated levels of CO_2_. Mineral analysis revealed that *N. khasiana* peristome and lid EFNs are rich in potassium, sodium, and calcium. The role of minerals in FNs and EFNs of *Nepenthes* is so far uninvestigated (Nicolson, 2022). Both *N. khasiana* EFNs showed high C:N ratios. Briefly, EFN secreted by *Nepenthes* is a sweet sugary mix, and due to minimal content of amino acids, proteins and vitamin C, it does not serve the nutritional requirements for visiting ants and other insects.

### AChE inhibition, active principle

In this study, the active principle, (+)-isoshinanolone, has been isolated from *N. khasiana* EFNs and its pitcher tissues. Several studies have demonstrated the occurrence of toxic secondary metabolites in FNs, and their role in favoring specialist pollinators and nectar protection (Adler, 2000; Heil, 2011; Stevenson *et al*., 2017; Barberis *et al*., 2023). These reports favor the proposition that the toxic principle is being transported from the phloem into the nectaries of *Nepenthes* pitchers.

Acetylcholine (ACh) is one of the key excitatory central nervous system neurotransmitters in vertebrates and invertebrates; and nerve impulse signal transmission at the synapses is accomplished through its degradation into choline and acetic acid by AChE (Colović *et al*., 2013). Our results showed that *N. khasiana* peristome and lid EFNs, their fractions and the active principle, (+)-isoshinanolone, inhibited AChE *in vitro*. Similarly, nectar-fed ants (their homogenates) collected from pitcher fluids (*in vivo*) proved significant inhibition to the kinetics of AChE. It is well established that ACh accumulation at the neuromuscular junctions causes symptoms of neurotoxicity such as cramps, muscular weakness, muscular fasciculation, paralysis, blurry vision, and even death (cholinergic toxicity) (Kalamida *et al*., 2007; Colović *et al*., 2013). Similarly, AChE inhibition by *Nepenthes* EFNs and their active constituent causes ACh accumulation, and consequent neurotoxic effects in ants and other visiting insects. These effects were further demonstrated by direct-feeding (*in vivo*) of EFN and the AChE inhibitor, (+)-isoshinanolone (IC_50_ 4.45 μg/ml), which resulted in cholinergic toxicity including death in ants (Pope *et al*., 2005; Colović *et al*., 2013; Supplementary Videos S1, S2). *N. khasiana* pitcher fluids with prey capture also showed AChE inhibition at high doses, but lower ingestible volumes by (fallen) ants (Paul and Roces, 2003) showed only negligible activity. EFN with its toxic effects affecting the mobility of visiting insects on a wet-low friction-slippery peristome (Bohn and Federle, 2004; Chen *et al*., 2016; Supplementary Video S4), with the waxy-slippery zone underneath and toxic viscoelastic digestive fluid (Gaume and Forterre, 2007) at the base of the pitcher are crucial designs favoring prey capture (Fig. 5). Ants are the predominant group of preys in *Nepenthes* (Fig. 1); *A. gracilipes* and *M. minutum* are the two major species of captured ants found seasonally in *N. khasiana* pitchers. Though *N. khasiana* EFN is sugar-rich, and therefore an energy source (Nicolson, 2022), it functions as a palatable ‘bait carrier’ against ants (Lee and Scotty Yang, 2022). Williamson and co-workers (2013) demonstrated cholinergic effects in honey bees (*Apis mellifera* Linn.) on treatment with AChE inhibitors. Moreover, Ratsirarson and Silander Jr. (1996) reported dizziness in insects on *N. madagascariensis* nectar consumption, enhancing the chances of them falling into the pitchers. Sluggish behavior and hampered flight of toxic nectar-fed pollinators were reported by various groups (Barberis *et al*., 2023 and references therein). Briefly, our study demonstrates (+)-isoshinanolone as a neuroactive compound which manipulates the physical behavior of insects. These results clearly establish *Nepenthes* EFN as a toxic sugar-rich secretion, which aids prey capture.

**Fig. 5.**
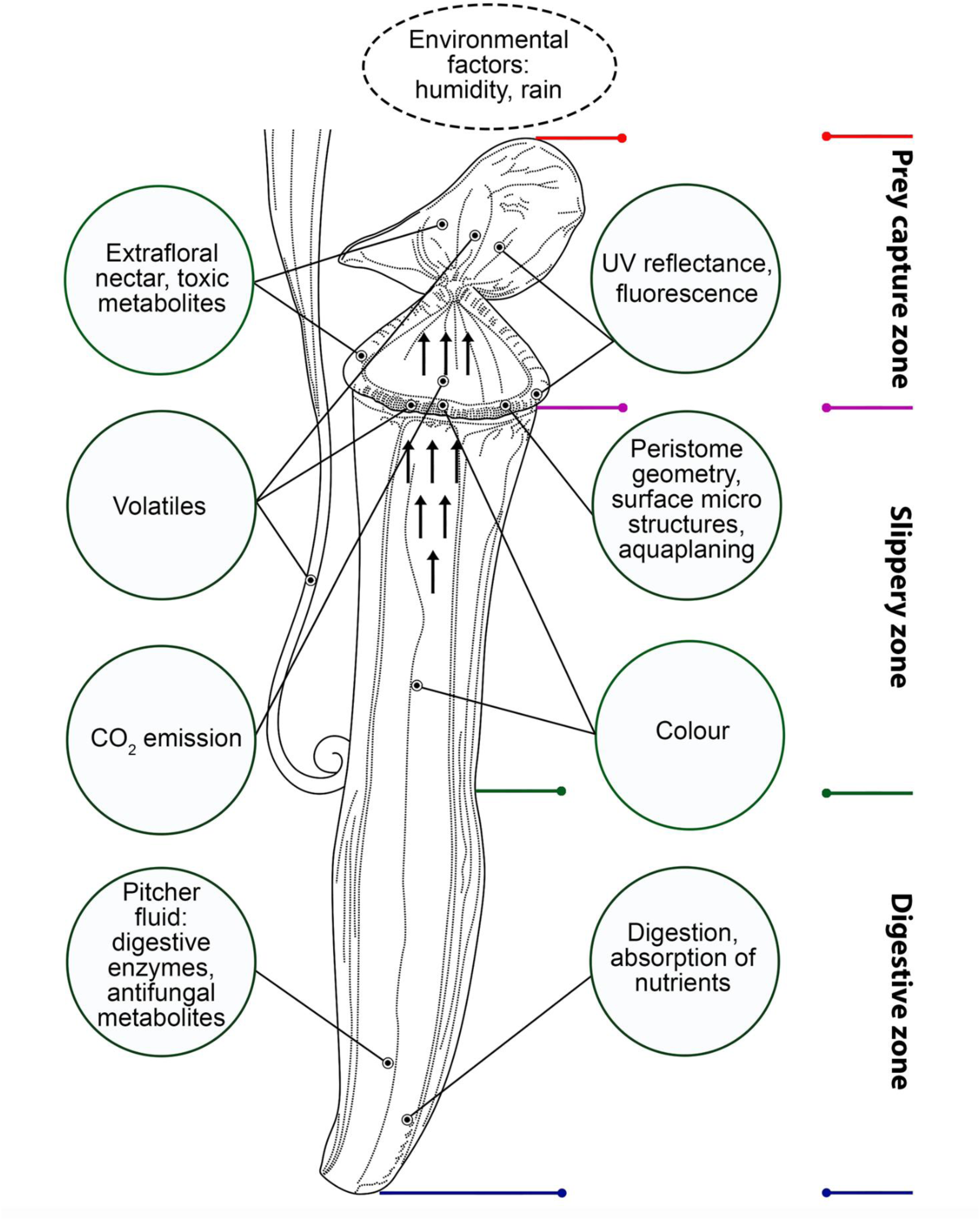
Prey capture strategies adopted by *Nepenthes* pitchers.

Even though *Nepenthes* pitchers are effective traps, and ants are their most captured preys (Fig. 1; Moran, 1996; Di Giusto *et al*., 2008; Gaume *et al*., 2016), there are factors denying the eventual capture of visiting ants by *Nepenthes* pitchers, *viz.*, toxic secondary metabolite-embedded nectar acts as a deterrent to visiting ants, and on assessing the palatability (toxicity), ants reject the nectar (Adler, 2000; Grasso *et al*., 2015; Stevenson *et al*., 2017). Besides, nectar ingestion causes uneven behavioral responses in ants. Considering these factors and other nutrient acquisition designs listed in Fig. 5, the extent of influence of EFN and its constituents in prey capture in *Nepenthes* is to be further investigated.

Bioactivity guided isolation of *N. khasiana* EFNs led to the AChE inhibitor, (+)-isoshinanolone, which is known for its fish-stunning and mosquito larvicidal activities. Similar to the EFNs, (+)-isoshinanolone showed green fluorescence at UV 366 nm (Hanson *et al*., 1981; Kurup *et al*., 2013). AChE inhibitors are known insecticides (Pope *et al*., 2005; Colović *et al*., 2013); therefore, (+)-isoshinanolone is a promising insecticidal entity. Moreover, it displayed AChE inhibition comparable to the Alzheimer’s disease (AD) drug, donepezil, and hence, it is a likely candidate for AD therapy (dos Santos *et al*., 2018). Isoshinanolone, an intermediate in plumbagin biosynthesis (Vasav *et al*., 2022), has been reported in several other *Nepenthes* species/hybrids (examples: *N. gracilis*, *N. thorelii*, *N. mirabilis*, *N. thorelii* x (*ventricosa* x *maxima*)) and other related genera (*Drosera indica*, *D. serpens*, *D. finlaysoniana*, *Dionaea*, *Drosophyllum*) (Hatcher *et al*., 2020; Wójciak *et al*., 2023). These observations prompt the chemical and toxicity evaluation of EFNs of other phylogenetically related carnivorous species.

### Volatiles, preys, herbivores

In our field study, ants were the most common prey type lured to *N. khasiana* pitchers (Fig. 1). This observation is in agreement with the previous studies in other native habitats, and the number of ants captured was positively correlated to EFN production (Moran, 1996; Merbach *et al*., 2001; Bauer *et al*., 2008; Di Giusto *et al*., 2008; Gaume *et al*., 2016). EFN is an easily accessible and relatively consistent food source for flightless ants. They are attracted to EFN-bearing tissues, and due to their presence plants generally benefit a reduction in herbivory; therefore, extrafloral nectaries and their associated nectar function as an indirect defense mechanism (Nicolson, 2007, 2022; Heil, 2015). We found plumbagin as a major volatile entity in the tendril and pitcher tissues in *khasiana*. Recent studies demonstrated the protection of *Nepenthes* against insect herbivores by naphthoquinones acting as phytoanticipins (Dávila-Lara *et al*., 2021b; Rahman-Soad *et al*., 2021). *Nepenthes khasiana* lid EFN on direct headspace-GC-MS analysis also showed traces of plumbagin, which is known for its antifeedant and antimicrobial activities (Raj *et al*., 2011). Plumbagin showed stronger AChE inhibition (IC_50_ 2.43 µg/ml) compared to (+)-isoshinanolone (IC_50_ 4.45 µg/ml) and the drug control, donepezil (IC_50_ 5.25 µg/ml). Both these naphthoquinones are found in *N. khasiana* tissues (Raj *et al*., 2011), and their neurotoxic effects possibly deterring herbivores on *Nepenthes* are to be ascertained (Grasso *et al*., 2015). *Nepenthes khasiana* EFNs showed very high C:N ratios. Its leaves (C 45.72 ± 2.43%, N 2.14 ± 0.30%, n = 4), and pitchers (C 39.07 ± 1.94%, N 1.50 ± 0.25%, n = 4) growing under the influence of high CO_2_ also displayed high C:N ratios (Osunkoya *et al*., 2007; Baby *et al*., 2017), which causes dilution of nitrogen resulting in reduced palatability and nutritional quality (Nicolson, 2022). Therefore, the reduction in herbivory observed in *Nepenthes* leaves and traps (Merbach *et al*., 2007; Mithöfer, 2022) is explained by ant visitation to these tissues, their nitrogen deficiency and high naphthoquinone contents.

### CO_2_, metabolism, prey capture

We have discovered 2500-5000 ppm of respiratory CO_2_ within unopened pitchers of *N. khasiana* and *Nepenthes* hybrids. Open *Nepenthes* pitchers emit CO_2_ as plumes above atmospheric levels (Baby *et al*., 2017). Studies demonstrated that CO_2_ plumes emitted by flowers act as indicators of adequate nectar supply to visiting pollinators (Goyret *et al*., 2008). In our study, CO_2_ streaming through *N. khasiana* open pitchers (620 ppm) caught a higher number of aerial preys compared to normal (CO_2_ 477 ppm) and air (CO_2_ 413 ppm)-streamed pitchers (Baby *et al*., 2017). Coincidentally, *Nepenthes* pitchers confront an elevated atmospheric CO_2_ scenario externally (419.3 ppm in 2023; Lindsey, 2024), in addition to the ten-times high CO_2_ within them (Baby *et al*., 2017).

Elevated CO_2_ induces changes in plant metabolism and enhances carbon assimilation demonstrated mainly by increased leaf C:N ratio and non-structural carbohydrates (Sardans *et al*., 2012; Zavala *et al*., 2013). *Nepenthes* follow the C3 photosynthetic pathway; and similar to the CO_2_-enrichment experiments, *N. khasiana* leaf and pitcher tissues showed high carbon and low nitrogen contents (Baby *et al*., 2017). Further, Osunkoya and co-workers (2007) recorded low nitrogen contents and high C:N ratios in leaves and pitchers of several *Nepenthes* species/hybrids from Brunei. High C:N ratio promotes the synthesis of C-based metabolites and limits the N-based ones. Again, high carbohydrate contents in the phloem tissues are transported and enzymatically transformed into nectar sugars in *Nepenthes* pitcher nectaries (Lin *et al*., 2014; Heil, 2015; Minami *et al*., 2021; Nicolson, 2022). *Nepenthes* pitchers due to low nitrogen, chlorophyll content and stomatal density, and replacement of chlorophyll-containing cells with digestive glands, display relatively low photosynthetic rates, and thereby low photosynthetic nitrogen-use efficiency. Moreover, these prey traps growing in elevated CO_2_ display reduced Rubisco activity (Pavlovič and Saganová, 2015). Pavlovič and Kocáb (2022) reported high level of alternative oxidase (AOX) in *Nepenthes* traps compared to their leaves, and inferred that part of the CO_2_ released from the pitchers is from the AOX activity. Another recent transcriptome sequencing study on the natural hybrid *Nepenthes x ventrata* ascertained the role of CO_2_ in pitcher development, function and evolution of its other key features (Shchennikova *et al*., 2021). CO_2_ within the pitchers also helps in maintaining the low pH of the digestive fluid (Baby *et al*., 2017).

Several secondary metabolites function as toxins in FNs (Adler, 2000; Grasso *et al*., 2015; Stevenson *et al*., 2017; Barberis *et al*., 2023). A few studies also mentioned anesthetizing alkaloids in *Nepenthes* nectar secretions (Bohn and Federle, 2004; Raj *et al*., 2011). But so far, there are no reports on the isolation of any nitrogenous secondary metabolites (alkaloids) in genus *Nepenthes*. Most secondary compounds reported from *Nepenthes* are naphthoquinones, and the second major group is phenolics (Hatcher *et al*., 2020; Wójciak *et al*., 2023). Naphthoquinones are considered as chemotaxonomic markers of the genus. Plumbagin and (+)-isoshinanolone, two major naphthoquinones, are proved to be neurotoxic to insects in this study. Two other naphthoquinones, droserone, 5-O-methyl droserone, were reported in *N. khasiana* pitcher fluid (Raj *et al*., 2011); they could be the entities causing fluid toxicity. Pitcher growth, high respiration, high carbohydrate, low nitrogenous primary and secondary metabolites, modified stomata and acidic pitcher fluid are critical features impelled by the high CO_2_ atmosphere within *Nepenthes* pitchers (Baby *et al*., 2017). In other words, *Nepenthes* pitchers are classic examples of leaf-evolved structures displaying manifestations of an elevated CO_2_ atmosphere within them.

Chemical (EFN, volatiles), gaseous (CO_2_), visual (colour, UV reflectance, fluorescence), physical (peristome geometry, surface microstructure, wettable surface-aquaplaning) and environmental (humidity, rain) factors are the prey capture strategies adopted by *Nepenthes* pitchers (Fig. 5). This study clearly demonstrates that *Nepenthes* EFNs provide only minimal nutritional benefits, instead they function as toxic baits aiding prey capture. In pollination, FN and pollen are true rewards, and both plant and pollinator mutually benefit, whereas in prey capture, the present discovery of a neurotoxin embedded in *Nepenthes* EFN nullifies the notion of reward to the visiting ants and other insects.

## Conclusions

Extrafloral nectar was believed to be the major reward in *Nepenthes* prey traps. This study demonstrated EFN secreted by *Nepenthes* pitchers as a sugar (Suc-Glc-Fru) mix, with low levels of key nutritional components, *viz.*, amino acids, proteins and vitamin C. Crucially, EFN is toxic to ants and other insects, and its toxic component (natural AChE inhibitor) has been identified as the naphthoquinone derivative, (+)-isoshinanolone. EFN secreted by *Nepenthes* unveils dual function of a lure and a toxin; and, unlike FNs, it provides only minimal nutritional benefits to the visitors. Our results prompt the chemical characterization and insect toxicity evaluation of naphthoquinones and EFNs in other *Nepenthes* species. The discovery of a toxic entity in EFN could also provide clues in better understanding the pollinator-prey conflict in *Nepenthes*. CO_2_ influenced metabolism, naphthoquinones (plumbagin, (+)-isoshinanolone) and ant visitation are the reasons for reduced herbivory in *Nepenthes*, particularly in their pitcher traps. High respiratory CO_2_ within *Nepenthes* pitchers results in their enhanced growth rate, C:N ratio, low Rubisco activity, modified stomata, acidic pitcher fluid and minimal nitrogenous components in its metabolism. Moreover, the multitudes of physico-chemical and environmental strategies of prey capture in *Nepenthes* are deceptive, and essentially no reward is being offered to the visiting ants and insects. Our findings infer elevated CO_2_ within their pitchers as the single most critical factor influencing the growth, metabolism, herbivory, and carnivory in *Nepenthes*.

## Supplementary data

The following supplementary data are available at JXB online. Details of the isolation of the active principle, (+)-isoshinanolone.

Table S1. Sugar composition in peristome and lid EFNs of *Nepenthes* species and hybrids.

Tabe S2. Mineral analysis of *N. khasiana* peristome and lid extrafloral nectars by ICP-OES.

Table S3. Insects and other organisms trapped in *N. khasiana* pitchers from 2 days to 14 days after opening.

Table S4. AChE inhibition by homogenates of captured ants (*in vivo*) and pitcher fluid collected from *N. khasiana* pitchers.

Table S5. Spectral data of (+)-isoshinanolone.

Table S6. AChE inhibition of *N. khasiana* EFNs and their active principle, (+)-isoshinanolone (*in vitro*).

Table S7. AChE inhibition of *Nepenthes* species, hybrids peristome and lid EFNs (*in vitro*).

Table S8. AChE inhibition by plumbagin.

Table S9. AChE inhibition by *N. khasiana* EFN– and (+)-isoshinanolone-fed ants (*A. gracilipes*, *in vivo*).

Fig. S1. Amino acid profiling in peristome and lid EFNs of *N. khasiana*.

Fig. S2. UFLC profiles of *N. khasiana* EFNs.

Fig. S3. ^1^H NMR of (+)-isoshinanolone.

Fig. S4. ^13^C NMR of (+)-isoshinanolone.

Fig. S5. Mass spectrum of (+)-isoshinanolone.

Fig. S6. Metabolic profiling of *N. khasiana* pitcher lid EFN by LC-MS.

Fig. S7. AChE inhibition of *Nepenthes* species peristome and lid EFNs (*in vitro*).

Fig. S8. AChE inhibition of *Nepenthes* hybrid peristome and lid EFNs (*in vitro*).

Fig. S9. AChE inhibition by plumbagin (*in vitro*).

Fig. S10. Detection of (+)-isoshinanolone in *Nepenthes* species/hybrid peristome and lid EFNs.

Video S1. Biotest: Behavioral pattern of *N. khasiana* peristome EFN-fed *A. gracilipes* ants (9 h).

Video S2. Biotest: Behavioral pattern of sugar mix + (+)-isoshinanolone-fed *A. gracilipes* ants (9 h).

Video S3. Biotest: Behavioral pattern of sugar mix (only)-fed *A. gracilipes* ants (9 h).

Video S4. Biotest: Behavioral pattern of control (water only-fed) *A. gracilipes* ants (9 h).

Video S5. Behavioral pattern of *A. gracilipes* ants on the peristome and leaves/tendrils of *N. khasiana*.

## Supporting information

003b_Nepenthes Supp Info_bioRxiv

## Acknowledgements

We are grateful to Dr. R. Raj Vikraman and Dr. K.J. Lathan Kumar, Garden Management Division, Jawaharlal Nehru Tropical Botanic Garden and Research Institute, Palode for providing us the plant specimens and facilities for field studies. We thank Mr. Manoj Vembayam, Travancore Natural History Society, Thiruvananthapuram, Kerala, India for identification of the ants. We acknowledge Dr. K. Madhavan Nampoothiri, CSIR-National Institute for Interdisciplinary Science and Technology, Thiruvananthapuram, Kerala, India for the UFLC analysis of amino acids. Authors thank Dr. K. C. Koshy, Fmr. Head, Plant Genetic Resource Division, Jawaharlal Nehru Tropical Botanic Garden and Research Institute for critical reading and suggestions on the manuscript.

## Author contributions

S.B., C.C.L., G.B.S., G.T., and A.J.J.: conceptualization; C.C.L., G.B.S., G.T., A.J.J., and S.B.: methodology, formal analysis and data acquisition; G.V.: biochemical assays, data interpretation; T.S.V.: data acquisition on insect visitors, interpretation, identification of ants; C.C.L., G.B.S., G.T., A.J.J., S.M., and L.A.S.: *Nepenthes* specimen collection, field support; S.B.: fund acquisition; S.B.: writing – original draft; S.B., C.C.L., G.B.S., A.J.J, G.V., and T.S.V: writing – review & editing; all authors contributed in finalizing the manuscript.

## Data availability

All data necessary to evaluate the conclusions of this paper are included in the article and/or supplementary data.

