## Supplementary material for "Bait, not reward: CO_2_-enriched *Nepenthes* pitchers secrete toxic nectar": 003b_Nepenthes Supp Info_bioRxiv

**Highlight:** *Nepenthes* extrafloral nectar has high content of carbohydrates and minimal nitrogenous metabolites. It is laced with (+)-isoshinanolone, an acetylcholinesterase inhibitor, and acts as a toxic bait, aiding prey capture.


### Abstract

*Nepenthes* pitchers are leaf-evolved biological traps holding high levels of $CO_2$ within them. Extrafloral nectar (EFN) secreted by these pitchers has long been regarded as the major reward to visiting arthropods, but its chemical constituents and their role in prey capture are least explored. Here we demonstrate *Nepenthes* EFN as a sugar (glucose-fructose-sucrose) mix with high C:N ratio, minimal amino acids, proteins, and vitamin C. *Nepenthes khasiana* peristome and lid EFNs displayed strong acetylcholinesterase (AChE) inhibition; the naphthoquinone derivative, (+)-isoshinanolone, has been identified as the AChE inhibitor. Plumbagin, the major volatile naphthoquinone in *Nepenthes*, also showed strong AChE inhibition. Direct EFN- and (+)-isoshinanolone-feeding bioassays demonstrated symptoms of cholinergic toxicity in ants. We testify that *Nepenthes* EFN is a toxic bait which hinders neuronal activity in visiting arthropods. These unique traps adopt various deceptive strategies for prey capture, and our discovery abolishes the notion that *Nepenthes* EFN is a reward to visiting ants and other arthropods. Moreover, our findings infer elevated $CO_2$ within their pitchers as the key factor influencing the growth, metabolism, herbivory, and carnivory in *Nepenthes*.



**Keywords:** C:N ratio, elevated $CO_2$, extrafloral nectar, (+)-isoshinanolone, naphthoquinone, nectar toxicity, *Nepenthes*, prey capture.

### Materials and methods

#### Activity guided isolation

Peristome (270.4 mg) and lid (249.1 mg) EFNs isolated from 48 *N. khasiana* pitchers were subjected to column chromatography separately over silica gel (200-400 mesh, 15.0 g, 1.8 x 60 cm), and the columns were eluted with hexane (100%), hexane:$CHCl_3$ (75:25, 50:50, 40:60, 30:70, 20:80, 10:90, 0:100%), $CHCl_3$:MeOH (95:5, 90:10, 75:25, 50:50) and MeOH (100%). Similar fractions were pooled together based on TLC profiles, which resulted in six major fractions from each column. Peristome EFN fractions: NPF1 (0.4 mg, 75:25 hexane:$CHCl_3$), NPF2 (0.7 mg, 50:50 hexane:$CHCl_3$), NPF3 (0.4 mg, 40:60 hexane:$CHCl_3$), NPF4 (1.1 mg, 20:80 hexane:$CHCl_3$), NPF5 (1.1 mg, 10:90 hexane:$CHCl_3$), NPF6 (188.4 mg, 75:25, 50:50 $CHCl_3$:MeOH, MeOH 100%). Lid EFN fractions: NLF1 (1.5 mg, 75:25 hexane:$CHCl_3$), NLF2 (0.1 mg, 50:50 hexane:$CHCl_3$), NLF3 (0.5 mg, 30:70 hexane:$CHCl_3$), NLF4 (1.0 mg, 10:90 hexane:$CHCl_3$), NLF5 (0.5 mg, $CHCl_3$ 100%), NLF6 (92.1 mg, 75:25, 50:50 $CHCl_3$:MeOH, MeOH 100%). These major fractions were tested for AChE inhibition, and NPF5 and NLF4 showed significant activity. TLC profiles of NPF5 and NLF4 displayed a major band at the same retention factor. To isolate this compound, peristome EFN (458.0 mg, isolated from 59 pitchers) was subjected to column chromatography over silica gel (200-400 mesh, 15 g, 1.8 x 60 cm), eluted with hexane (100%), hexane:$CHCl_3$ (75:25, 50:50, 40:60, 30:70, 20:80, 10:90), $CHCl_3$ (100%), $CHCl_3$:MeOH (95:5, 90:10, 75:25, 50:50) and MeOH (100%). Six major fractions were obtained, of which fifth fraction yielded the active component (3.1 mg only).

To obtain higher amounts of the active principle, *N. khasiana* peristome water extract was tested for this component, and co-TLC confirmed its presence in it. Peristomes of 30 *N. khasiana* pitchers were separated (18.6 g), ground to small pieces, extracted with distilled water (100 ml) using a magnetic stirrer (150 rpm, 3 h) and the filtered extract was evaporated to dryness using a rotary evaporator at 37$^{o}$C under reduced pressure. This extract (1.2 g) was subjected to column chromatography over silica gel (200-400 mesh, 36.0 g, 4.5 x 60 cm), eluting with hexane (100%), hexane:$CHCl_3$ (75:25, 50:50, 40:60, 30:70, 20:80, 10:90), $CHCl_3$ (100%), $CHCl_3$:MeOH (95:5, 90:10, 75:25, 50:50) and MeOH

(100%). Similar fractions were pooled together, and the impure active compound (10.0 mg) was obtained from the sixth fraction (20:80 hexane:$CHCl_3$). These extraction-isolation steps were repeated three times to obtain 23.5 mg of the impure fraction. Re-chromatography of this fraction over silica gel (200-400 mesh, 10 g, 1.8 x 45 cm) by eluting with hexane:$CHCl_3$ (40:60, 30:70, 20:80, 10:90) yielded the active compound (14.0 mg; 20:80 hexane:$CHCl_3$). This isolation protocol was repeated to obtain sufficient quantity of the active principle for the experiments.

**Table S1. Sugar composition in peristome and lid EFNs of *Nepenthes* species and hybrids.**

| *Nepenthes* species/ hybrid | Pitcher length (cm) | Peristome diameter (cm) | Lid length (cm) | Digestive zone diameter (cm) | Pitcher fluid (ml) | Fresh weight | | Nectar yield (mg) | Suc (%)‡ | Glu (%)‡ | Fru (%)‡ |
|---|---|---|---|---|---|---|---|---|---|---|---|
| | | | | | | Peristome (g) | Lid (g) | | | | |
| *N. khasiana* peristome | | | | | | | | | | | |
| P1 | 20.2 | 3.9 | 6.0 | 3.5 | 10.0 | 1.12 | 1.03 | 7.0 | 6.92 ± 0.18 | 7.53 ± 0.32 | 10.06 ± 0.37 |
| P2 | 22.3 | 4.5 | 6.5 | 4.3 | 15.0 | 1.54 | 1.50 | 13.2 | 3.42 ± 0.26 | 4.75 ± 0.28 | 9.14 ± 0.27 |
| P3 | 18.3 | 3.3 | 5.3 | 2.8 | 10.0 | 0.87 | 0.91 | 4.2 | 6.20 ± 0.13 | 6.81 ± 0.17 | 9.41 ± 0.31 |
| P4 | 16.7 | 3.4 | 4.9 | 3.3 | 7.0 | 0.76 | 0.74 | 5.3 | 6.86 ± 0.36 | 7.16 ± 0.15 | 11.49 ± 0.68 |
| P5 | 25.6 | 5.2 | 7.4 | 4.5 | 17.0 | 2.20 | 2.13 | 25.9 | 9.78 ± 0.24 | 9.90 ± 0.30 | 21.49 ± 0.35 |
| P6 | 17.2 | 3.5 | 5.4 | 3.5 | 10.0 | 0.92 | 0.88 | 4.4 | ND (0.00) | ND (0.00) | ND (0.00) |
| Average | | | | | | | | 10.0 ± 8.48 (n = 6) | 5.53 ± 3.38% (n = 6) | 6.03 ± 3.38% (n = 6) | 10.27 ± 6.85% (n = 6) |
| *N. khasiana* lid | | | | | | | | | | | |
| L1 | 20.2 | 3.9 | 6.0 | 3.5 | 10.0 | 1.12 | 1.03 | 16.2 | 8.42 ± 0.62 | 16.24 ± 0.82 | 22.35 ± 0.84 |
| L2 | 22.3 | 4.5 | 6.5 | 4.3 | 15.0 | 1.54 | 1.50 | 4.1 | ND (0.00) | 8.63 ± 0.31 | 22.19 ± 0.30 |
| L3 | 18.3 | 3.3 | 5.3 | 2.8 | 10.0 | 0.87 | 0.91 | 4.1 | 6.65 ± 0.38 | 9.98 ± 0.63 | 17.18 ± 0.45 |

| | | | | | | | | | | | |
|---|---|---|---|---|---|---|---|---|---|---|---|
| L4 | 16.7 | 3.4 | 4.9 | 3.3 | 7.0 | 0.76 | 0.74 | 3.5 | 2.58 ± 0.00 | 5.32 ± 0.82 | 14.78 ± 1.13 |
| L5 | 25.6 | 5.2 | 7.4 | 4.5 | 17.0 | 2.20 | 2.13 | 5.7 | 4.39 ± 0.60 | 6.41 ± 0.84 | 15.03 ± 2.09 |
| L6 | 17.2 | 3.5 | 5.4 | 3.5 | 10.0 | 0.92 | 0.88 | 3.8 | ND (0.00) | ND (0.00) | 3.24 ± 0.27 |
| Average | | | | | | | | 6.23 ± 4.94 (n = 6) | 3.67 ± 3.47% (n = 6) | 7.76 ± 5.40% (n = 6) | 15.80 ± 7.00 (n = 6) |
| *N. mirabilis* peristome | | | | | | | | | | | |
| P1 | 9.4 | 2.0 | 2.5 | 2.6 | 3.0 | 0.22 | 0.08 | 1.6 | ND (0.00) | ND (0.00) | ND (0.00) |
| P2 | 7.2 | 1.6 | 2.9 | 2.9 | 7.6 | 0.23 | 0.08 | 2.1 | 2.29 ± 0.28 | ND (0.00) | ND (0.00) |
| Average | | | | | | | | 1.85 ± 0.35 (n = 2) | 1.15 ± 1.62 (n = 2) | 0.00 ± 0.00 (n = 2) | 0.00 ± 0.00 (n = 2) |
| *N. mirabilis* lid | | | | | | | | | | | |
| L1 | 9.4 | 2.0 | 2.5 | 2.6 | 3.0 | 0.22 | 0.08 | 1.8 | ND (0.00) | ND (0.00) | ND (0.00) |
| L2 | 7.2 | 1.6 | 2.9 | 2.9 | 7.6 | 0.23 | 0.08 | 2.1 | 7.40 ± 0.07 | ND (0.00) | ND (0.00) |
| Average | | | | | | | | 1.95 ± 0.21 (n = 2) | 3.70 ± 5.23 (n = 2) | 0.00 ± 0.00 (n = 2) | 0.00 ± 0.00 (n = 2) |
| *N. mirabilis* × *N. khasiana* peristome | | | | | | | | | | | |
| P1 | 16.7 | 3.3 | 4.0 | 3.2 | 11.0 | 0.36 | 0.23 | 16.9 | 14.69 ± 0.43 | 4.81 ± 0.38 | 5.32 ± 0.39 |

| | | | | | | | | | | | |
|---|---|---|---|---|---|---|---|---|---|---|---|
| P2 | 15.2 | 4.3 | 4.6 | 3.9 | 19.0 | 0.41 | 0.23 | 1.5 | 20.68 ± 0.34 | ND (0.00) | ND (0.00) |
| P3 | 18.2 | 4.1 | 4.3 | 3.6 | 23.0 | 0.53 | 0.31 | 4.4 | 4.98 ± 0.06 | 6.25 ± 0.25 | 5.60 ± 0.22 |
| P4 | 18.5 | 4.0 | 5.0 | 3.2 | 20.0 | 0.70 | 0.39 | 8.1 | 8.21 ± 0.13 | 6.07 ± 0.22 | 9.85 ± 0.23 |
| P5 | 18.1 | 4.5 | 4.2 | 3.2 | 12.0 | 0.92 | 0.46 | 10.8 | 27.11 ± 0.31 | 3.68 ± 0.12 | 4.28 ± 0.19 |
| Average | | | | | | | | 8.34 ± 5.95 (n = 5) | 15.13 ± 9.02 (n = 5) | 4.16 ± 2.55 (n = 5) | 5.01 ± 3.52 (n = 5) |
| *N. mirabilis* × *N. khasiana* lid | | | | | | | | | | | |
| L1 | 16.7 | 3.3 | 4.0 | 3.2 | 11.0 | 0.36 | 0.23 | 4.3 | 24.56 ± 0.48 | 13.44 ± 0.33 | 16.10 ± 0.29 |
| L2 | 15.2 | 4.3 | 4.6 | 3.9 | 19.0 | 0.41 | 0.23 | 0.3 | ND (0.00) | ND (0.00) | ND (0.00) |
| L3 | 18.2 | 4.1 | 4.3 | 3.6 | 23.0 | 0.53 | 0.31 | 1.3 | 1.01 ± 2.03 | ND (0.00) | 0.85 ± 1.70 |
| L4 | 18.5 | 4.0 | 5.0 | 3.2 | 20.0 | 0.70 | 0.39 | 0.2 | ND (0.00) | ND (0.00) | ND (0.00) |
| L5 | 18.1 | 4.5 | 4.2 | 3.2 | 12.0 | 0.92 | 0.46 | 8.4 | 35.08 ± 0.61 | 3.85 ± 0.22 | 3.56 ± 0.11 |
| Average | | | | | | | | 2.90 ± 3.49 (n = 5 | 12.13 ± 16.58 (n = 5) | 3.46 ± 5.82 (n = 5) | 4.10 ± 6.86 (n = 5) |
| *N. mirabilis* × *N. rafflesiana* peristome | | | | | | | | | | | |
| P1 | 19.6 | 4.7 | 4.9 | 2.3 | 17.5 | 0.99 | 0.27 | 13.7 | ND (0.00) | 3.19 ± 0.13 | 6.34 ± 0.20 |

| | | | | | | | | | | | |
|---|---|---|---|---|---|---|---|---|---|---|---|
| P2 | 19.0 | 5.2 | 4.9 | 1.6 | 14.0 | 1.43 | 0.50 | 11.3 | 0.11 ± 0.01 | 2.35 ± 0.32 | 3.69 ± 0.30 |
| P3 | 16.6 | 3.9 | 3.8 | 1.9 | 30.0 | 0.59 | 0.15 | 3.9 | 2.67 ± 0.17 | ND (0.00) | ND (0.00) |
| Average | | | | | | | | 9.63 ± 5.11 (n = 3) | 0.93 ± 1.51 (n = 3) | 1.85 ± 1.65 (n = 3) | 3.34 ± 3.18 (n = 3) |
| *N. mirabilis* × *N. rafflesiana* lid | | | | | | | | | | | |
| L1 | 19.6 | 4.7 | 4.9 | 2.3 | 17.5 | 0.99 | 0.27 | 3.4 | ND (0.00) | ND (0.00) | ND (0.00) |
| L2 | 19.0 | 5.2 | 4.9 | 1.6 | 14.0 | 1.43 | 0.50 | 8.0 | 9.56 ± 0.39 | ND (0.00) | ND (0.00) |
| L3 | 16.6 | 3.9 | 3.8 | 1.9 | 30.0 | 0.59 | 0.15 | 5.1 | ND (0.00) | ND (0.00) | ND (0.00) |
| Average | | | | | | | | 5.5 ± 2.33 (n = 3) | 3.19 ± 5.52 (n = 3) | 0.00 ± 0.00 (n = 3) | 0.00 ± 0.00 (n = 3) |

‡Each  Suc, Glu, Fru % is an average of four data points;  ND: Not detected.

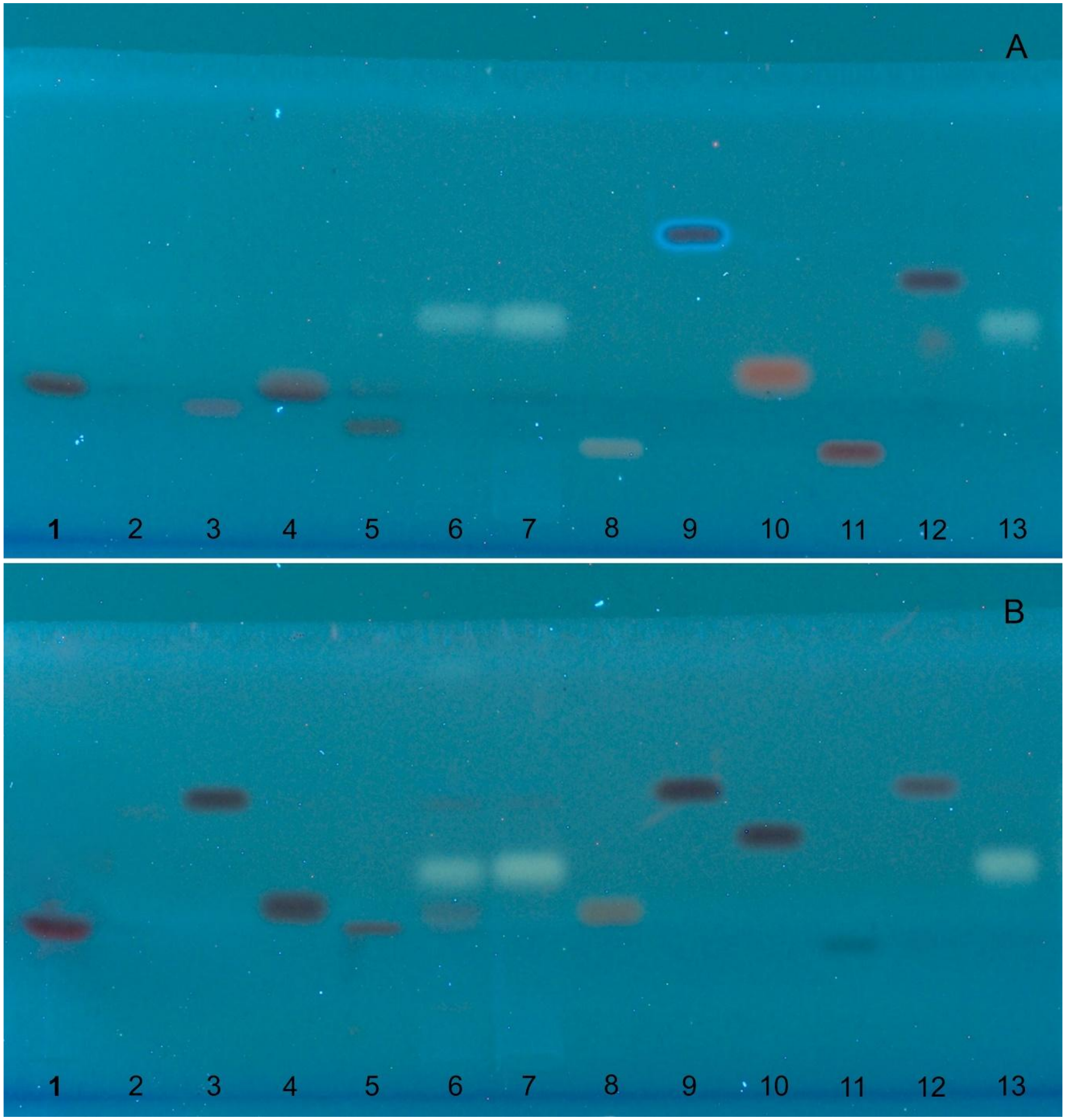
A
1 2 3 4 5 6 7 8 9 10 11 12 13
B
1 2 3 4 5 6 7 8 9 10 11 12 13

**Fig. S1. Amino acid profiling in peristome and lid EFNs of *N. khasiana*.** (**A**): Track 1: Gln, track 2: Cys, track 3: Asp, track 4: Gly, track 5: Arg, track 6: *NKP-EFN, track 7: **NKL-EFN, track 8: His, track 9: Trp, track 10: Pro, track 11: Lys, track 12: Met, track 13: #Sugar (Suc-Glu-Fru) mix; (**B**): track 1: Ser, track 2: Tyr, track 3: Ile, track 4: Ala, track 5: Glu, track 6: *NKP-EFN, track 7: **NKL-EFN, track 8: Thr, track 9: Leu, track 10: Val, track 11: Asn, track 12: Phe, track 13: #Sugar (Suc-Glu-Fru) mix. All 20 standard amino acids were applied at 2 µg per track; *NKP-EFN & **NKL-EFN: *N. khasiana* peristome and lid EFNs (10 µg per track); #Sugar mix: (Suc:Glu:Fru 2.08:2.08:2.08 µg, 1.5 µl per track). HPTLC plates developed in butanol:acetic acid:water (12:3:5, v/v, 20 ml), derivatized using ninhydrin reagent, scanned at 570 nm and photographed at UV 366 nm in CAMAG TLC visualizer 2.0 (details in *Material and Methods, Amino acids*).

*N. khasiana* peristome and lid EFNs, even at 10 µg per track, did not show bands with Rf values matching to the 20 standard AAs (2 µg per track). Instead, EFNs showed unresolved sugar bands (matching the standard sugar mix in tracks 13) in these chromatographic conditions. Repeated analysis showed the same chromatographic profiles for *N. khasiana* peristome/lid EFNs. Moreover, EFNs of *Nepenthes* species/hybrids in this study gave negative results in ninhydrin tests for AAs.

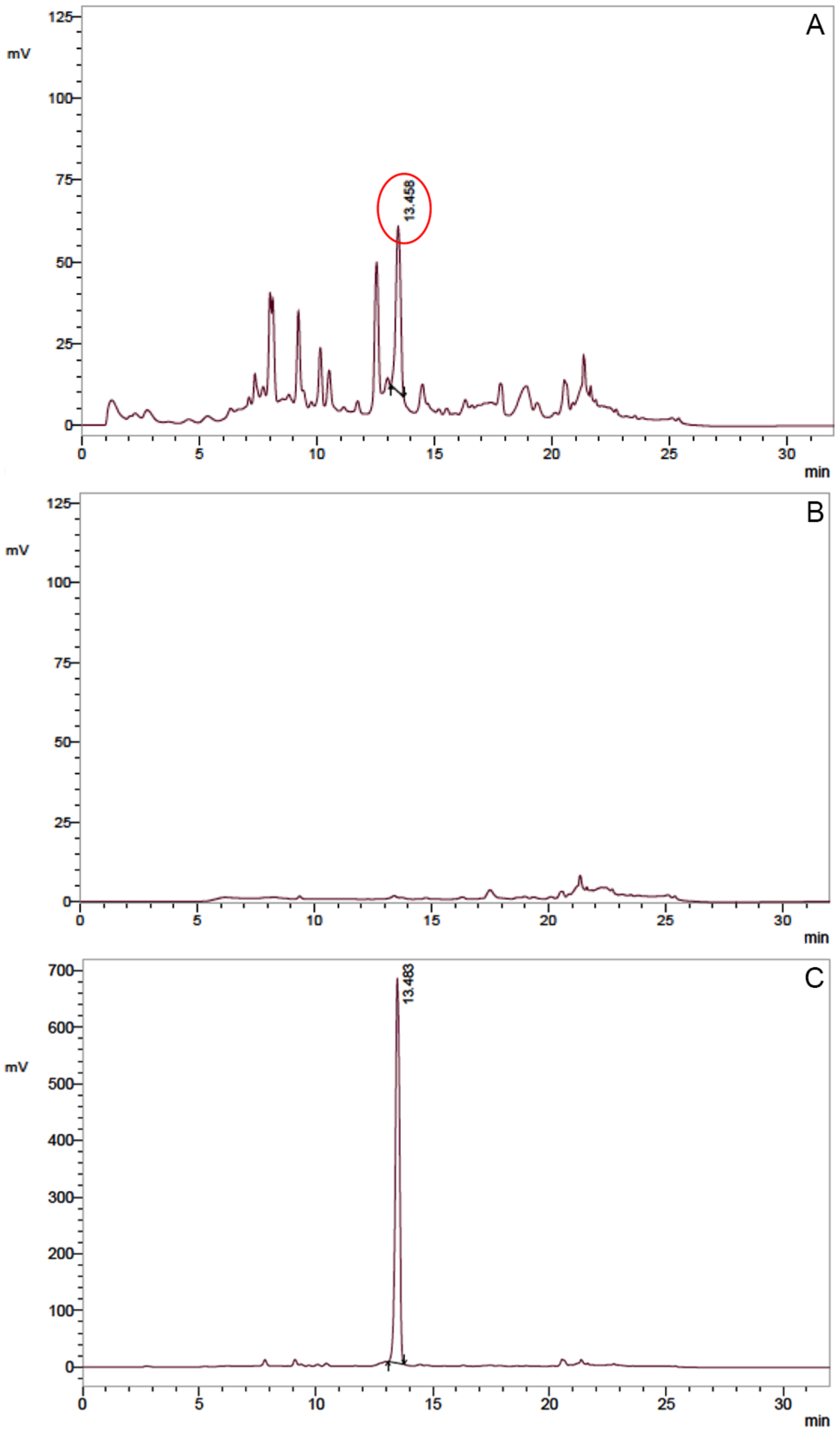


**Fig. S2. UFLC profiles of *N. khasiana* EFNs.** (A) peristome EFN, (B) lid EFN, and (C) methionine standard (2 µg/ml).

Amino acids in EFNs were also analysed using a Shimadzu UFLC system equipped with a fluorescence detector (RF 20A) at excitation and emission wavelengths of 348 and 450 nm, respectively. EFNs dissolved in Milli-Q water (peristome EFN: 11.4 mg/ml, lid EFN: 12.0 mg/ml; injection volume 10 µl each) were filtered through 0.22 mm filter membranes (PALL Life Sciences), buffered at pH 10.0 using borate buffer (Agilent) and pre-derivatized with o-phthaldialdehyde (OPA, Agilent Life Sciences). Amino acids were resolved with Zorbax Eclipse AAA column (5 µm, 4.6 × 150 mm; Agilent Technologies) using 40 mM sodium phosphate buffer (Agilent) and acetonitrile-methanol-water in the ratio (45:45:10) in a gradient elution program (Sasikumar *et al.*, 2021).

Of the 20 amino acid standards, only traces of methionine (0.0008%) was detected in peristome EFN; none of the amino acids were detected in the lid EFN.

**Table S2. Mineral analysis of *N. khasiana* peristome and lid extrafloral nectars by ICP-OES.**

| Element | Peristome EFN (mg/l) | Lid EFN (mg/l) |
|---|---|---|
| Ag | 0.23 ± 0.03 | 0.22 ± 0.02 |
| Al | 1.40 ± 0.28 | 1.58 ± 0.28 |
| B | 0.04 ± 0.01 | 0.43 ± 0.08 |
| Ba | BDL | BDL |
| Bi | 3.06 ± 0.61 | 3.03 ± 0.58 |
| Ca | 7.49 ± 1.18 | 23.42 ± 3.40 |
| Cd | BDL | BDL |
| Co | BDL | BDL |
| Cr | BDL | BDL |
| Cu | BDL | BDL |
| Fe | BDL | BDL |
| Ga | 1.12 ± 0.11 | 1.13 ± 0.17 |
| In | BDL | BDL |
| K | 48.14 ± 8.97 | 159.8 ± 31.96 |
| Mg | 3.83 ± 0.77 | 12.98 ± 2.59 |
| Mn | 0.58 ± 0.05 | BDL |
| Na | 24.21 ± 4.84 | 68.67 ± 13.74 |
| Ni | BDL | BDL |
| Pb | BDL | BDL |
| Tl | 0.53 ± 0.08 | 0.58 ± 0.09 |
| Zn | 0.78 ± 0.15 | 0.98 ± 0.20 |

Samples were analyzed in triplicate; results are mean ± S.D.; BDL: below detection limit; 1 mg/l = 1 ppm .

**Table S3. Insects and other organisms trapped in *N. khasiana* pitchers from 2 days to 14 days after opening.**

| Prey diversity | 2 DAO | 4 DAO | 6 DAO | 8 DAO | 10 DAO | 12 DAO | 14 DAO |
|---|---|---|---|---|---|---|---|
| Hymenoptera (other than ants) | 0.50 ± 0.84 | 0.50 ± 0.84 | 4.83 ± 7.08 | 1.50 ± 1.38 | 1.67 ± 1.63 | 6.67 ± 7.55 | 3.33 ± 3.67 |
| Ants | 500.00 ± 348.68 | 560.0 ± 556.05 | 1095.00 ± 874.70 | 2208.33 ± 2103.33 | 818.3 ± 649.53 | 457.5 ± 442.21 | 593.3 ± 619.61 |
| Lepidoptera | 0.33 ± 0.82 | 1.67 ± 1.37 | 5.84 ± 2.78 | 8.17 ± 8.33 | 3.50 ± 2.89 | 8.00 ± 7.23 | 10.00 ± 7.87 |
| Diptera | 2.84 ± 3.82 | 11.50 ± 10.38 | 28.67 ± 11.94 | 34.17 ± 19.69 | 25.17 ± 22.31 | 26.17 ± 31.51 | 29.00 + 19.14 |
| Coleoptera | 0.00 ± 0.00 | 0.50 ± 1.22 | 0.33 ± 0.52 | 0.33 ± 0.82 | 1.17 ± 0.75 | 0.33 ± 0.52 | 0.50 ± 0.84 |
| Hemiptera | 0.17 ± 0.41 | 0.33 ± 0.82 | 0.17 ± 0.41 | 0.00 ± 0.00 | 0.16 ± 0.41 | 0.00 ± 0.00 | 0.17 ± 0.41 |
| Dictyoptera | 0.00 ± 0.00 | 0.00 ± 0.00 | 0.00 ± 0.00 | 0.17 ± 0.41 | 0.50 ± 0.84 | 0.33 ± 0.52 | 0.00 ± 0.00 |
| Orthoptera | 0.00 ± 0.00 | 0.00 ± 0.00 | 0.33 ± 0.82 | 0.00 ± 0.00 | 0.00 ± 0.00 | 0.17 ± 0.41 | 0.50 ± 0.84 |
| Isoptera | 0.17 ± 0.41 | 0.00 ± 0.00 | 0.00 ± 0.00 | 0.00 ± 0.00 | 0.00 ± 0.00 | 0.00 ± 0.00 | 0.00 ± 0.00 |
| Neuroptera | 0.00 ± 0.00 | 0.00 ± 0.00 | 0.00 ± 0.00 | 0.00 ± 0.00 | 0.00 ± 0.00 | 0.17 ± 0.41 | 0.00 ± 0.00 |
| Thysanoptera | 0.33 ± 0.82 | 0.17 ± 0.41 | 0.00 ± 0.00 | 0.00 ± 0.00 | 0.00 ± 0.00 | 0.00 ± 0.00 | 0.00 ± 0.00 |
| Arachnids (Spiders) | 0.17 ± 0.41 | 0.17 ± 0.41 | 0.00 ± 0.00 | 0.17 ± 0.41 | 1.83 ± 4.02 | 0.00 ± 0.00 | 0.67 ± 1.21 |
| Molluscs | 0.00 ± 0.00 | 0.00 ± 0.00 | 0.00 ± 0.00 | 0.00 ± 0.00 | 0.00 ± 0.00 | 0.00 ± 0.00 | 0.17 ± 0.41 |
| Reptiles | 0.00 ± 0.00 | 0.00 ± 0.00 | 0.00 ± 0.00 | 0.17 ± 0.41 | 0.00 ± 0.00 | 0.00 ± 0.00 | 0.00 ± 0.00 |
| Millipedes | 0.00 ± 0.00 | 0.00 ± 0.00 | 0.00 ± 0.00 | 0.00 ± 0.00 | 0.00 ± 0.00 | 0.00 ± 0.00 | 0.50 ± 1.22 |
| Maggots (Dipteran) | 0.00 ± 0.00 | 0.00 ± 0.00 | 0.00 ± 0.00 | 0.17 ± 0.41 | 0.00 ± 0.00 | 0.00 ± 0.00 | 3.83 ± 9.39 |

Sample collection: May to July 2018; data presented as average prey capture per pitcher ± S.D., n = 6; DAO: days after pitcher opening; *N. khasiana* pitchers start disintegrating after 14 days of prey capture; total prey capture: 38807; ants only: 37395 (96.4%). Sampling in November 2023 showed similar trends, with very high capture rate (%) of ants. These data are in agreement with previous reports from other natural habitats (Moran, 1996; Di Giusto *et al.*, 2008; Gaume *et al.*, 2016).

**Table S4. AChE inhibition by homogenates of captured ants (*in vivo*) and pitcher fluid collected from *N. khasiana* pitchers.**

| Sl. No. | Test material | Inhibition (%) |
|---|---|---|
| 1. | AChI (136 µg/ml) + AChE (5 units/l) | 1.0 ± 1.0 |
| 2. | AChI (136 µg/ml) + donepezil (2.5 µg/ml) + AChE (5 units/l) | 38.5 ± 1.5 |
| 3. | AChI (136 µg/ml) + donepezil (5.0 µg/ml) + AChE (5 units/l) | 53.0 ± 3.0 |
| 4. | AChI (136 µg/ml) + donepezil (10 µg/ml) + AChE (5 units/l) | 73.0 ± 4.0 |
| 5. | AChI (136 µg/ml) + donepezil (100 µg/ml) + AChE (5 units/l) | 87.0 ± 1.0 |
| 6. | AChI (136 µg/ml) + donepezil (1000 µg/ml) + AChE (5 units/l) | 91.0 ± 1.0 |
| 7. | AChI (136 µg/ml) + control-enzyme source (non-EFN fed ant *M. minutum* homogenate) (5 units/l) | 11.3 ± 1.2 |
| 8. | AChI (136 µg/ml) + test-enzyme source (EFN-fed ant *M. minutum* homogenate) (5 units/l) | 99.9 ± 0.1 |
| 9. | AChI (136 µg/ml) + control-enzyme source (non-EFN fed ant *A. gracilipes* homogenate) (5 units/l) | 14.0 ± 2.7 |
| 10. | AChI (136 µg/ml) + test-enzyme source (EFN-fed ant *A. gracilipes* homogenate) (5 units/l) | 99.9 ± 0.0 |
| 11. | AChI (136 µg/ml) + pitcher fluid (1 µl) + AChE (5 units/l) | 0.9 ± 0.2 |
| 12. | AChI (136 µg/ml) + pitcher fluid (2.5 µl) + AChE (5 units/l) | 1.4 ± 0.2 |
| 13. | AChI (136 µg/ml) + pitcher fluid (5 µl) + AChE (5 units/l) | 3.6 ± 0.4 |
| 14. | AChI (136 µg/ml) + pitcher fluid (10 µl) + AChE (5 units/l) | 4.6 ± 0.4 |
| 15. | AChI (136 µg/ml) + pitcher fluid (25 µl) + AChE (5 units/l) | 19.7 ± 2.5 |
| 16. | AChI (136 µg/ml) + pitcher fluid (50 µl) + AChE (5 units/l) | 59.3 ± 3.1 |
| 17. | AChI (136 µg/ml) + pitcher fluid (100 µl) + AChE (5 units/l) | 94.7 ± 1.5 |

AChI (substrate): 100 µl; donepezil (drug): 100 µl; AChE: 50 µl; ant homogenates: 50 µl each; pitcher fluid is the digestive fluid from *N. khasiana* pitcher with prey capture; samples were assayed in triplicate.

**Table S5. Spectral data of (+)-isoshinanolone** (Hanson *et al.*, 1981)

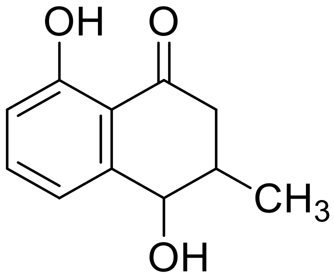

(+)-Isoshinanolone

$[\alpha]_D^{22} = +30.0°$ ($CHCl_3$, c. 0.97%)

IR $v_{max}$ (KBr) $cm^{-1}$: 3301 (O-H str.), 2925 (C-H str), 2910, 1659 (C=O str), 1430, 1030.

$^1$H NMR ($CDCl_3$, 400 MHz) δ (ppm): 2.56 (2H, m) H-2, 2.80 (1H, q) H-3, 4.71 (1H, dd) H-4, 3.44 (1H, s; OH), 6.9 (1H, d) H-5, 7.1 (1H, dd) H-6, 7.45 (1H, d) H-7, 12.39 (1H, s; OH), 1.15 (3H, s) H-11.

$^{13}$C NMR ($CDCl_3$, 100 MHz) δ (ppm): 205.16 (C-1), 40.72 (C-2), 34.43 (C-3), 71.11 (C-4), 118.09 (C-5), 145.09 (C-6), 118.71 (C-7), 162.67 (C-8), 11.93 (C-9), 136.95 (C-10), 16.16 (C-11).

HRMS: $[M+H]^+$ *m/z* 193.0862, calculated mass 192.0786.

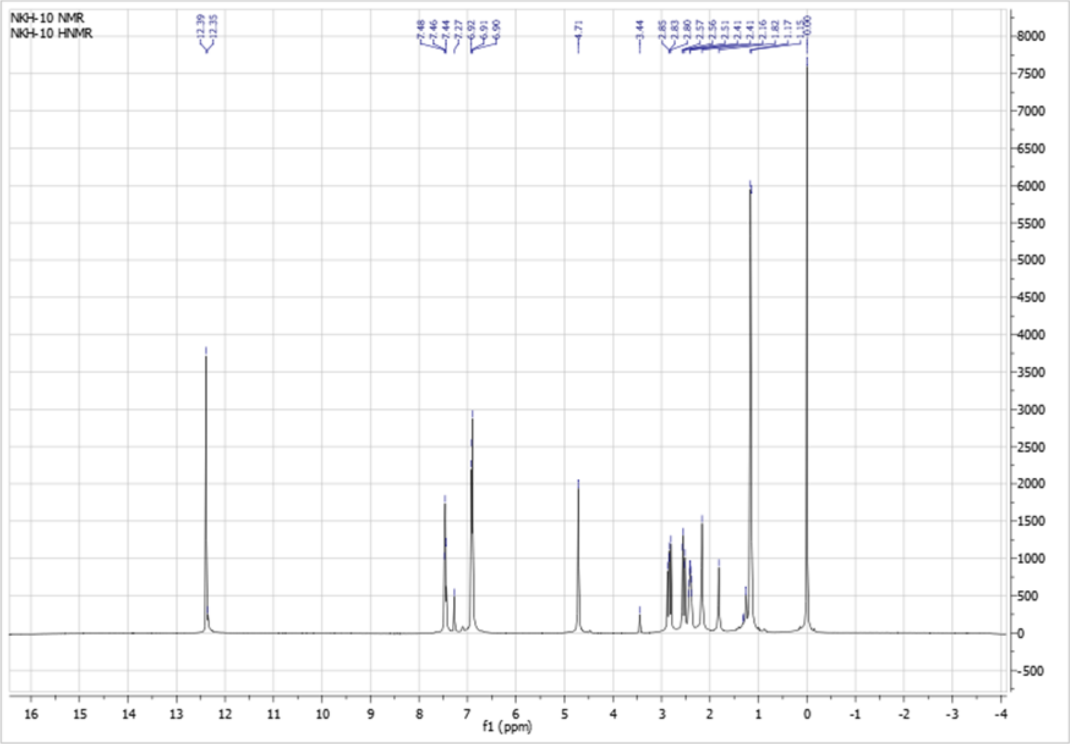


**Fig. S3.** $^{1}$H NMR of (+)-isoshinanolone.

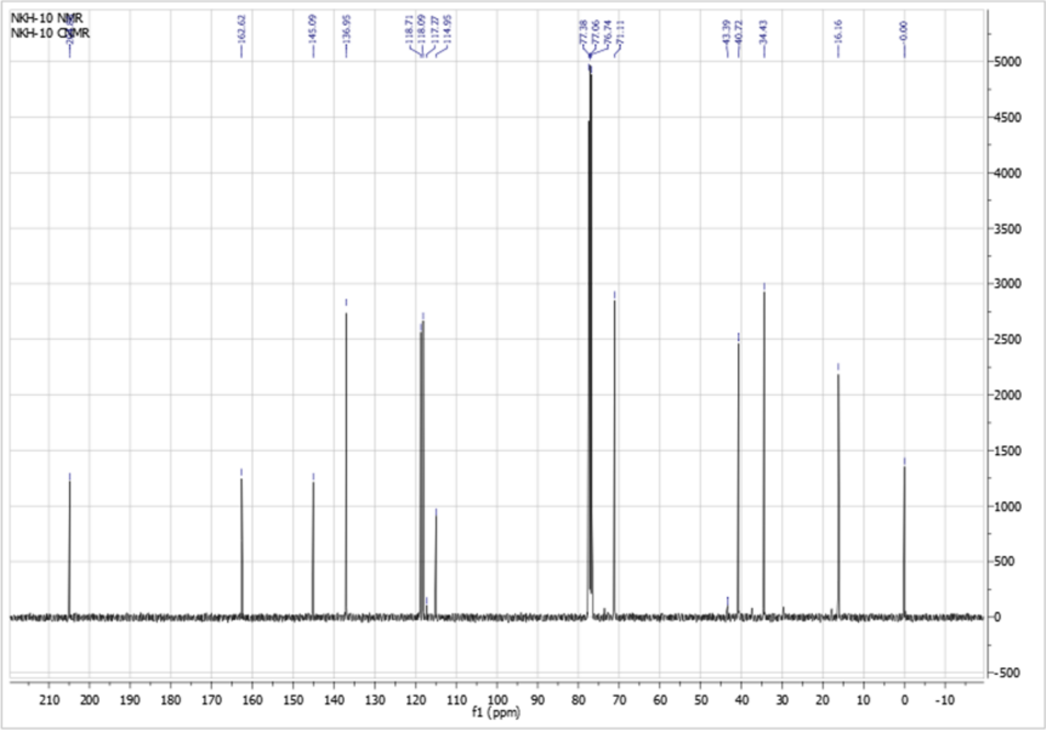


**Fig. S4.** $^{13}$C NMR of (+)-isoshinanolone.

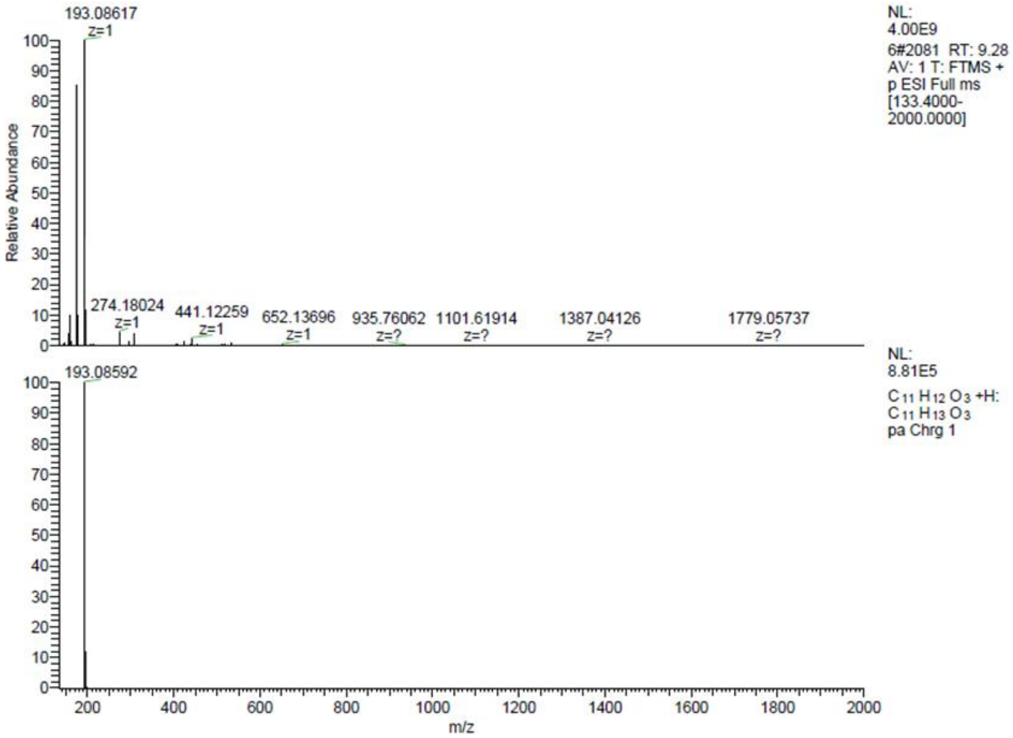


**Fig. S5**. Mass spectrum of (+)-isoshinanolone.

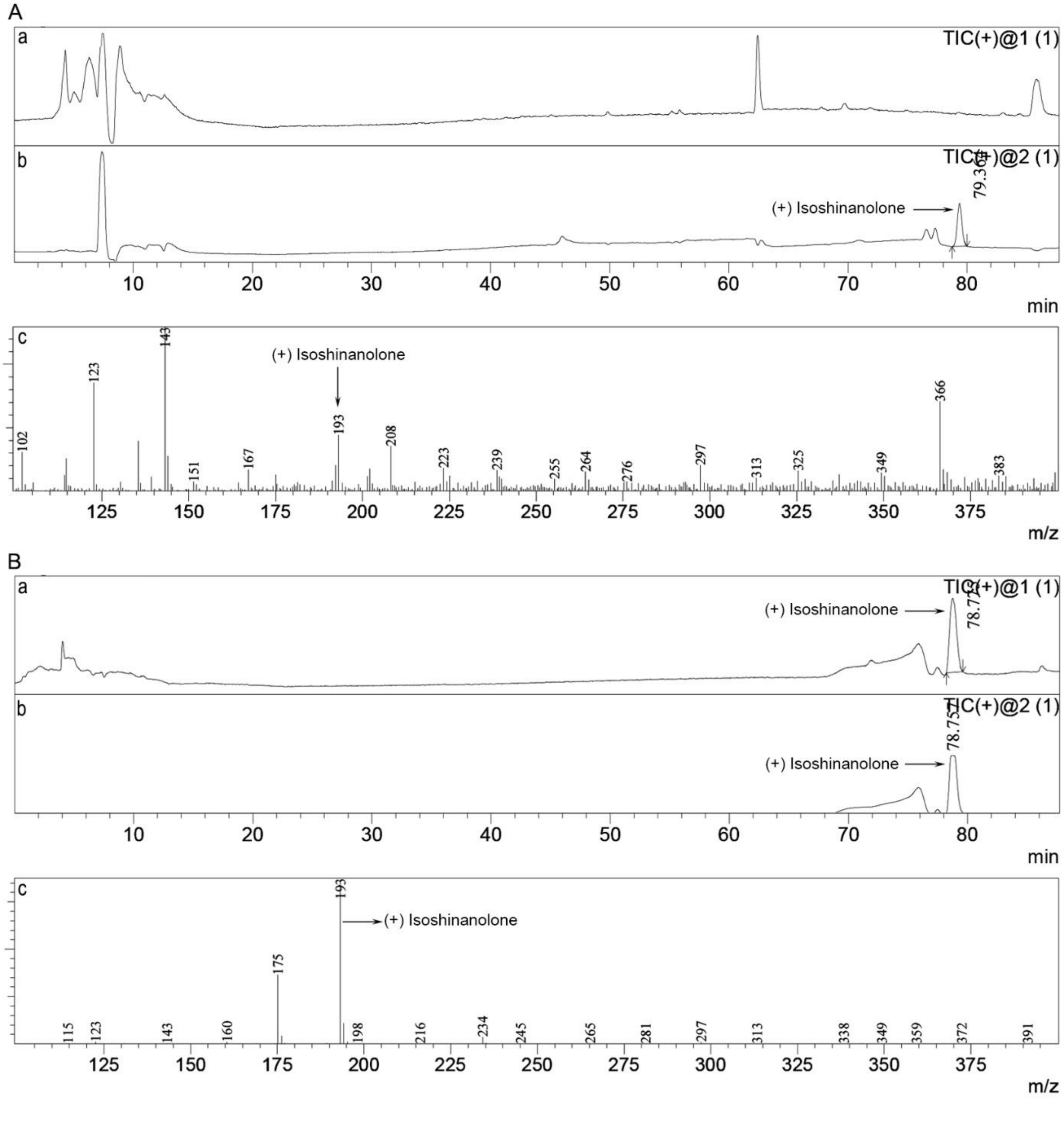


**Fig. S6. Metabolic profiling of *N. khasiana* pitcher lid EFN by LC-MS. A**. (a) TIC of *N. khasiana* lid nectar, (b) selective ion mode of lid nectar showing (+) isoshinanolone, (c) mass spectrum of lid nectar; **B**. (a) TIC of (+) isoshinanolone standard, (b) selective ion mode of (+) isoshinanolone, (c) mass spectrum of (+) isoshinanolone; Y-axes: relative abundance (in varying scales).

**Table S6. AChE inhibition of *N. khasiana* EFNs and their active principle, (+)-isoshinanolone (*in vitro*).**

| Sl. No. | Test material | Inhibition (%) |
|---|---|---|
| 1. | AChI (136 µg/ml) + AChE (5 units/l) | 3.4 ± 0.2 |
| 2. | AChI (136 µg/ml) + donepezil (1000 µg/ml) + AChE (5 units/l) | 91.0 ± 1.0 |
| 3. | AChI (136 µg/ml) + peristome EFN (1000 µg/ml) + AChE (5 units/l) | 87.0 ± 2.0 |
| 4. | AChI (136 µg/ml) + lid EFN (1000 µg/ml) + AChE (5 units/l) | 96.0 ± 1.0 |
| 5. | AChI (136 µg/ml) + (+)-isoshinanolone (10 µg/ml) + AChE (5 units/l) | 94.5 ± 0.5 |
| 6. | AChI (136 µg/ml) + (+)-isoshinanolone (100 µg/ml) + AChE (5 units/l) | 97.5 ± 0.5 |
| 7. | AChI (136 µg/ml) + (+)-isoshinanolone (1000 µg/ml) + AChE (5 units/l) | 100.0 ± 0.0 |

AChI (substrate): 100 µl; donepezil (drug): 100 µl; (+)-isoshinanolone: 100 µl; peristome and lid EFNs: 100 µl each; AChE: 50 µl; samples were assayed in triplicate.

**Table S7. AChE inhibition of *Nepenthes* species, hybrids peristome and lid EFNs (*in vitro*).**

| Sl. No. | Test material | Inhibition (%) |
|---|---|---|
| 1. | AChI (136 µg/ml) + AChE (5 units/l) | 3.4 ± 0.2 |
| 2. | AChI (136 µg/ml) + donepezil (1000 µg/ml) + AChE (5 units/l) | 91.0 ± 1.0 |
| 3. | AChI (136 µg/ml) + *N. mirabilis* peristome EFN (10 µg/ml) + AChE (5 units/l) | 55.3 ± 3.3 |
| 4. | AChI (136 µg/ml) + *N. mirabilis* peristome EFN (100 µg/ml) + AChE (5 units/l) | 62.3 ± 3.3 |
| 5. | AChI (136 µg/ml) + *N. mirabilis* peristome EFN (1000 µg/ml) + AChE (5 units/l) | 77.9 ± 1.9 |
| 6. | AChI (136 µg/ml) + *N. mirabilis* lid EFN (10 µg/ml) + AChE (5 units/l) | 50.3 ± 1.7 |
| 7. | AChI (136 µg/ml) + *N. mirabilis* lid EFN (100 µg/ml) + AChE (5 units/l) | 60.7 ± 3.3 |
| 8. | AChI (136 µg/ml) + *N. mirabilis* lid EFN (1000 µg/ml) + AChE (5 units/l) | 74.7 ± 3.3 |
| 9. | AChI (136 µg/ml) + *N. khasiana* peristome EFN (10 µg/ml) + AChE (5 units/l) | 52.5 ± 2.5 |
| 10. | AChI (136 µg/ml) +*N. khasiana* peristome EFN (100 µg/ml) + AChE (5 units/l) | 75.0 ± 2.0 |
| 11. | AChI (136 µg/ml) +*N. khasiana* peristome EFN (1000 µg/ml) + AChE (5 units/l) | 87.0 ± 2.0 |
| 12. | AChI (136 µg/ml) + *N. khasiana* lid EFN (10 µg/ml) + AChE (5 units/l) | 62.5 ± 2.5 |
| 13. | AChI (136 µg/ml) + *N. khasiana* lid EFN (100 µg/ml) + AChE (5 units/l) | 83.5 ± 1.5 |
| 14. | AChI (136 µg/ml) + *N. khasiana* lid EFN (1000 µg/ml) + AChE (5 units/l) | 96.0 ± 1.0 |

| | | |
|---|---|---|
| 15. | AChI (136 µg/ml) + *N. mirabilis* × *N. khasiana* peristome EFN (10 µg/ml) + AChE (5 units/l) | 38.2 ± 1.9 |
| 16. | AChI (136 µg/ml) + *N. mirabilis* × *N. khasiana* peristome EFN (100 µg/ml) + AChE (5 units/l) | 41.7 ± 0.0 |
| 17. | AChI (136 µg/ml) + *N. mirabilis* × *N. khasiana* peristome EFN (1000 µg/ml) + AChE (5 units/l) | 52.3 ± 1.8 |
| 18. | AChI (136 µg/ml) + *N. mirabilis* × *N. khasiana* lid EFN (10 µg/ml) + AChE (5 units/l) | 32.9 ± 1.9 |
| 19. | AChI (136 µg/ml) + *N. mirabilis* × *N. khasiana* lid EFN (100 µg/ml) + AChE (5 units/l) | 40.7 ± 0.0 |
| 20. | AChI (136 µg/ml) + *N. mirabilis* × *N. khasiana* lid EFN (1000 µg/ml) + AChE (5 units/l) | 43.7 ± 0.1 |
| 21. | AChI (136 µg/ml) + *N. mirabilis* × *N. rafflesiana* peristome EFN (10 µg/ml) + AChE (5 units/l) | 40.3 ± 0.1 |
| 22. | AChI (136 µg/ml) + *N. mirabilis* × *N. rafflesiana* peristome EFN (100 µg/ml) + AChE (5 units/l) | 44.0 ± 0.1 |
| 23. | AChI (136 µg/ml) + *N. mirabilis* × *N. rafflesiana* peristome EFN (1000 µg/ml) + AChE (5 units/l) | 46.0 ± 0.1 |
| 24. | AChI (136 µg/ml) + *N. mirabilis* × *N. rafflesiana* lid EFN (10 µg/ml) + AChE (5 units/l) | 34.0 ± 0.1 |
| 25. | AChI (136 µg/ml) + *N. mirabilis* × *N. rafflesiana* lid EFN (100 µg/ml) + AChE (5 units/l) | 39.3 ± 0.0 |
| 26. | AChI (136 µg/ml) + *N. mirabilis* × *N. rafflesiana* lid EFN (1000 µg/ml) + AChE (5 units/l) | 53.0 ± 0.0 |

AChI (substrate): 100 µl; donepezil (drug): 100 µl; peristome and lid EFNs: 100 µl each; AChE: 50 µl; samples were assayed in triplicate.

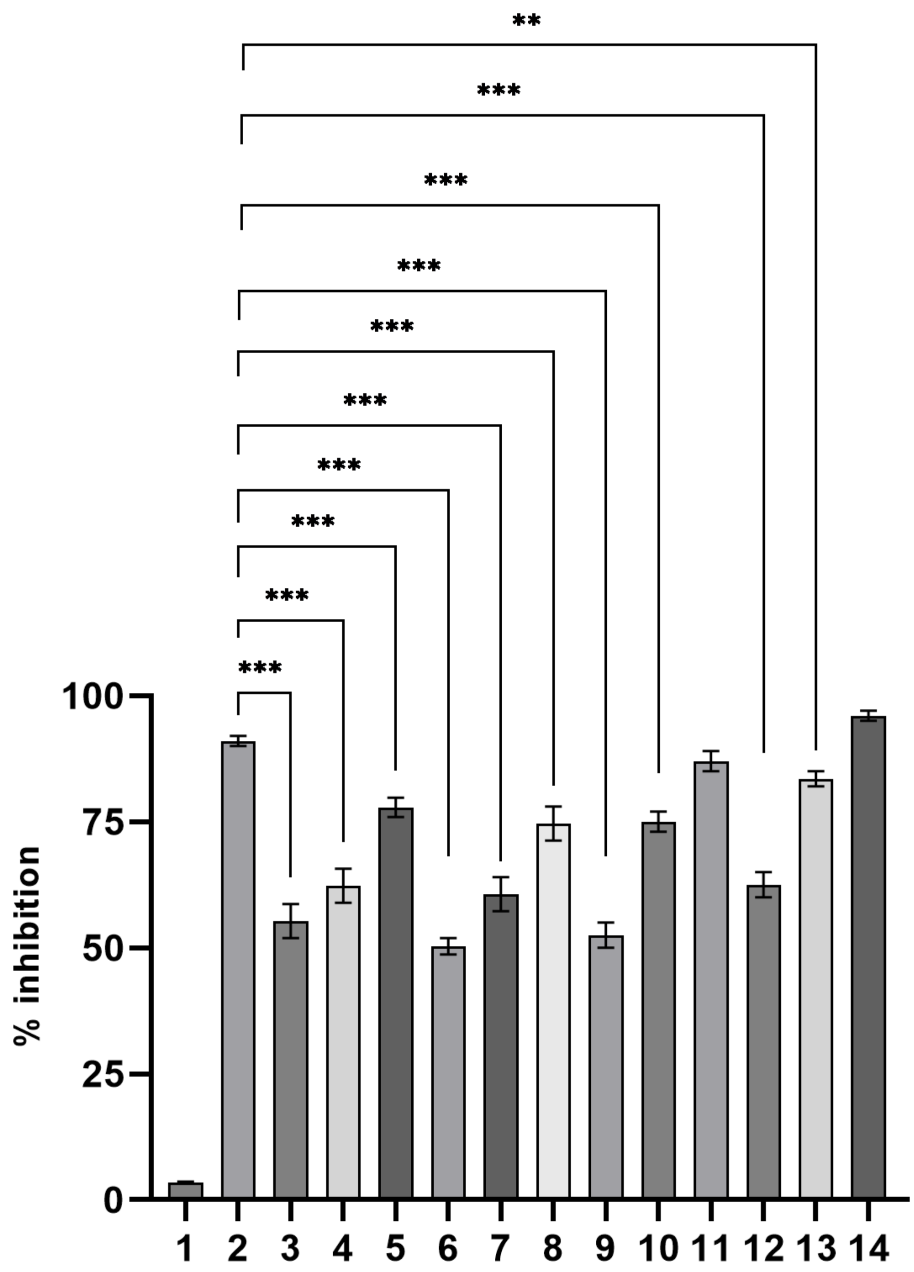


**Fig. S7. AChE inhibition of *Nepenthes* species peristome and lid EFNs (*in vitro*).** (**1**: AChI (136 µg/ml) + AChE (5 units/l); **2**: AChI (136 µg/ml) + donepezil (1000 µg/ml) + AChE (5 units/l); **3**: AChI (136 µg/ml) + *N. mirabilis* peristome EFN (10 µg/ml) + AChE (5 units/l); **4**: AChI (136 µg/ml) + *N. mirabilis* peristome EFN (100 µg/ml) + AChE (5 units/l); **5**: AChI (136 µg/ml) + *N. mirabilis* peristome EFN (1000 µg/ml) + AChE (5 units/l); **6**: AChI (136 µg/ml) + *N. mirabilis* lid EFN (10 µg/ml) + AChE (5 units/l); **7**: AChI (136 µg/ml) + *N. mirabilis* lid EFN (100 µg/ml) + AChE (5 units/l); **8**: AChI (136 µg/ml) + *N. mirabilis* lid EFN (1000 µg/ml) + AChE (5 units/l); **9**: AChI (136 µg/ml) + *N. khasiana* peristome EFN (10 µg/ml + AChE (5 units/l); **10**: AChI (136 µg/ml) +*N. khasiana* peristome EFN (100 µg/ml) + AChE (5 units/l); **11**: AChI (136 µg/ml) +*N. khasiana* peristome EFN (1000 µg/ml) + AChE (5 units/l); **12**: AChI (136 µg/ml) + *N. khasiana* lid EFN (10 µg/ml) + AChE (5 units/l); **13**: AChI (136 µg/ml) + *N. khasiana* lid EFN (100 µg/ml) + AChE (5 units/l); **14**: AChI (136 µg/ml) + *N. khasiana* lid EFN (1000 µg/ml) + AChE (5 units/l)). $p$ values were calculated using one-way ANOVA and Sidak's multiple comparison test using GraphPad Prism 8.0.2; values are mean ± S.D.; n = 3; $^{**}p = 0.0062$; $^{***}p < 0.0001$ (compared to 2); data listed in Table S7.

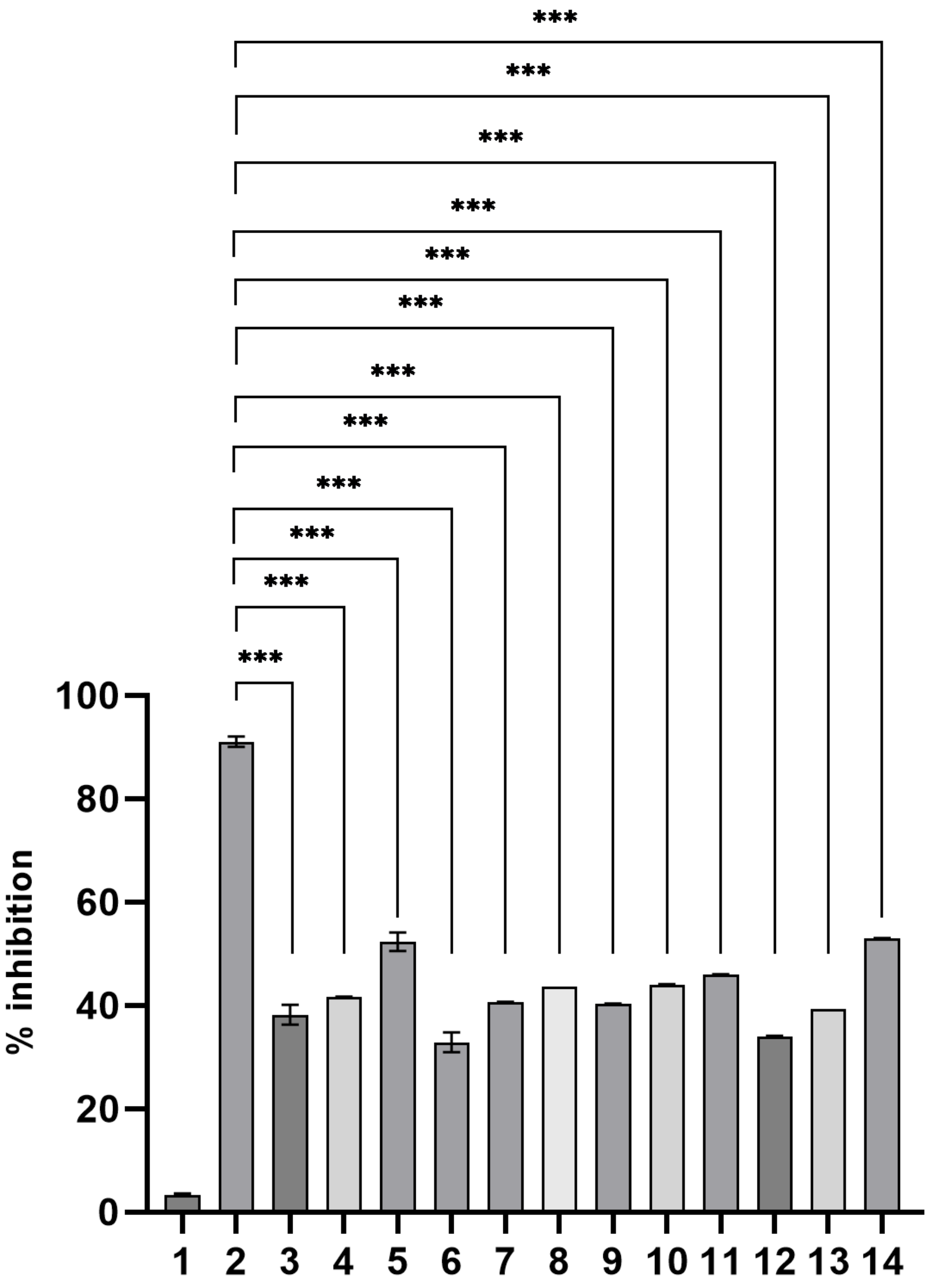


**Fig. S8. AChE inhibition of *Nepenthes* hybrid peristome and lid EFNs (*in vitro*).** (**1**: AChI (136 µg/ml) + AChE (5 units/l); **2**: AChI (136 µg/ml) + donepezil (1000 µg/ml) + AChE (5 units/l); **3**: AChI (136 µg/ml) + *N. mirabilis* × *N. khasiana* peristome EFN (10 µg/ml) + AChE (5 units/l); **4**: AChI (136 µg/ml) + *N. mirabilis* × *N. khasiana* peristome EFN (100 µg/ml) + AChE (5 units/l); **5**: AChI (136 µg/ml) + *N. mirabilis* × *N. khasiana* peristome EFN (1000 µg/ml) + AChE (5 units/l); **6**: AChI (136 µg/ml) + *N. mirabilis* × *N. khasiana* lid EFN (10 µg/ml) + AChE (5 units/l); **7**: AChI (136 µg/ml) + *N. mirabilis* × *N. khasiana* lid EFN (100 µg/ml) + AChE (5 units/l); **8**: AChI (136 µg/ml) + *N. mirabilis* × *N. khasiana* lid EFN

(1000 µg/ml) + AChE (5 units/l); **9**: AChI (136 µg/ml) + *N. mirabilis* × *N. rafflesiana* peristome EFN (10 µg/ml) + AChE (5 units/l); **10**: AChI (136 µg/ml) + *N. mirabilis* × *N. rafflesiana* peristome EFN (100 µg/ml) + AChE (5 units/l); **11**: AChI (136 µg/ml) + *N. mirabilis* × *N. rafflesiana* peristome EFN (1000 µg/ml) + AChE (5 units/l); **12**: AChI (136 µg/ml) + *N. mirabilis* × *N. rafflesiana* lid EFN (10 µg/ml) + AChE (5 units/l); **13**: AChI (136 µg/ml) + *N. mirabilis* × *N. rafflesiana* lid EFN (100 µg/ml) + AChE (5 units/l); **14**: AChI (136 µg/ml) + *N. mirabilis* × *N. rafflesiana* lid EFN (1000 µg/ml) + AChE (5 units/l)). $p$ values were calculated using one-way ANOVA and Sidak's multiple comparison test using GraphPad Prism 8.0.2; values are mean ± S.D.; n = 3; ***$p < 0.0001$ (compared to 2); data listed in Table S7.

**Table S8. AChE inhibition by plumbagin.**

| Sl. No. | Test material | Inhibition (%) |
|---|---|---|
| 1. | AChl (136 µg/ml) + AChE (5 units/l) | 5.7 ± 3.1 |
| 2. | AChl (136 µg/ml) + donepezil (1000 µg/ml) + AChE (5 units/l) | 92.7 ± 5.2 |
| 3. | AChl (136 µg/ml) + plumbagin (1.25 µg/ml) + AChE (5 units/l) | 49.0 ± 6.8 |
| 4. | AChl (136 µg/ml) + plumbagin (2.5 µg/ml) + AChE (5 units/l) | 50.7 ± 4.9 |
| 5. | AChl (136 µg/ml) + plumbagin (5.0 µg/ml) + AChE (5 units/l) | 72.7 ± 9.5 |
| 6. | AChl (136 µg/ml) + plumbagin (10 µg/ml) + AChE (5 units/l) | 89.7 ± 1.2 |
| 7. | AChl (136 µg/ml) + plumbagin (100 µg/ml) + AChE (5 units/l) | 93.3 ± 1.2 |
| 8. | AChl (136 µg/ml) + plumbagin (1000 µg/ml) + AChE (5 units/l) | 98.3 ± 1.2 |

(AChl (substrate): 100 µl; donepezil (drug): 100 µl; plumbagin: 100 µl; AChE: 50 µl).

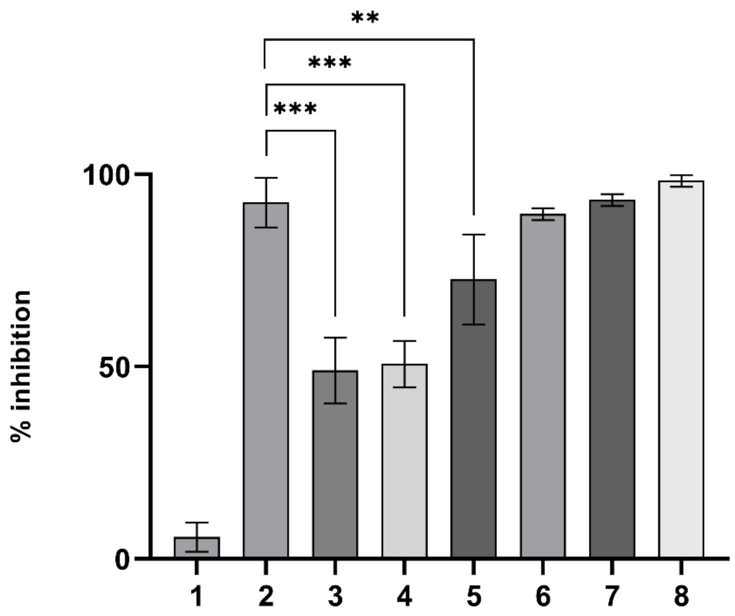


**Fig. S9. AChE inhibition by plumbagin (*in vitro*).** 1: AChl (136 µg/ml) + AChE (5 units/l); 2: AChl (136 µg/ml) + donepezil (1000 µg/ml) + AChE (5 units/l); 3: AChl (136 µg/ml) + plumbagin (1.25 µg/ml) + AChE (5 units/l); 4: AChl (136 µg/ml) + plumbagin (2.5 µg/ml) + AChE (5 units/l); 5: AChl (136 µg/ml) + plumbagin (5.0 µg/ml) + AChE (5 units/l); 6: AChl (136 µg/ml) + plumbagin (10 µg/ml) + AChE (5 units/l); 7: AChl (136 µg/ml) + plumbagin (100 µg/ml) + AChE (5 units/l); 8: AChl (136 µg/ml) + plumbagin (1000 µg/ml) + AChE (5 units/l)). *p* values were calculated using one-way ANOVA and Dunnett's multiple comparison test using GraphPad Prism 8.0.2; values are mean ± S.D.; n = 3; ***$p < 0.0001$ (compared to 2); **$p = 0.0065$; data listed in Table S8.

**Table S9. AChE inhibition by *N. khasiana* EFN- and (+)-isoshinanolone-fed ants (*A. gracilipes*, *in vivo*).**

| Sl. No. | Test material | Inhibition (%) |
|---|---|---|
| 1. | AChI (136 µg/ml) + unfed ant homogenate (5 units/l) | 4.6 ± 0.3 |
| 2. | AChI (136 µg/ml) + sugar mix (Suc 0.55 mg + Glc 0.60 mg + Fru 1.03 mg)-fed control ant homogenate (5 units/l) | 3.7 ± 0.1 |
| 3. | AChI (136 µg/ml) + (sugar mix + donepezil 1.25 µg/ml)-fed ant homogenate (5 units/l) | 26.0 ± 1.0 |
| 4. | AChI (136 µg/ml) + (sugar mix + donepezil 2.5 µg/ml)-fed ant homogenate (5 units/l) | 33.5 ± 0.5 |
| 5. | AChI (136 µg/ml) + (sugar mix + donepezil 5.0 µg/ml)-fed ant homogenate (5 units/l) | 48.5 ± 0.5 |
| 6. | AChI (136 µg/ml) + (sugar mix + donepezil 10 µg/ml)-fed ant homogenate (5 units/l) | 87.0 ± 1.0 |
| 7. | AChI (136 µg/ml) + (sugar mix + (+)-isoshinanolone 1.25 µg/ml)-fed ant homogenate (5 units/l) | 22.5 ± 0.5 |
| 8. | AChI (136 µg/ml) + (sugar mix + (+)-isoshinanolone 2.5 µg/ml)-fed ant homogenate (5 units/l) | 46.0 ± 1.0 |
| 9. | AChI (136 µg/ml) + (sugar mix + (+)-isoshinanolone 5.0 µg/ml)-fed ant homogenate (5 units/l) | 56.5 ± 1.5 |
| 10. | AChI (136 µg/ml) + (sugar mix + (+)-isoshinanolone 10.0 µg/ml)-fed ant homogenate (5 units/l) | 98.0 ± 1.0 |
| 11. | AChI (136 µg/ml) + *N. khasiana* peristome EFN (10 µg/ml)-fed ant homogenate (5 units/l) | 50.0 ± 1.0 |
| 12. | AChI (136 µg/ml) + *N. khasiana* peristome EFN (100 µg/ml)-fed ant homogenate (5 units/l) | 71.7 ± 1.5 |
| 13. | AChI (136 µg/ml) + *N. khasiana* peristome EFN (1000 µg/ml)-fed ant homogenate (5 units/l) | 86.0 ± 1.7 |
| 14. | AChI (136 µg/ml) + *N. khasiana* lid EFN (10 µg/ml)-fed ant homogenate (5 units/l) | 59.2 ± 1.0 |

| | | |
|---|---|---|
| 15. | AChl (136 µg/ml) + *N. khasiana* lid EFN (100 µg/ml)-fed ant homogenate (5 units/l) | 83.3 ± 1.3 |
| 16. | AChl (136 µg/ml) + *N. khasiana* lid EFN (1000 µg/ml)-fed ant homogenate (5 units/l) | 94.8 ± 0.8 |

AChl (substrate): 100 µl; ant homogenates: 50 µl each; samples were assayed in triplicate.

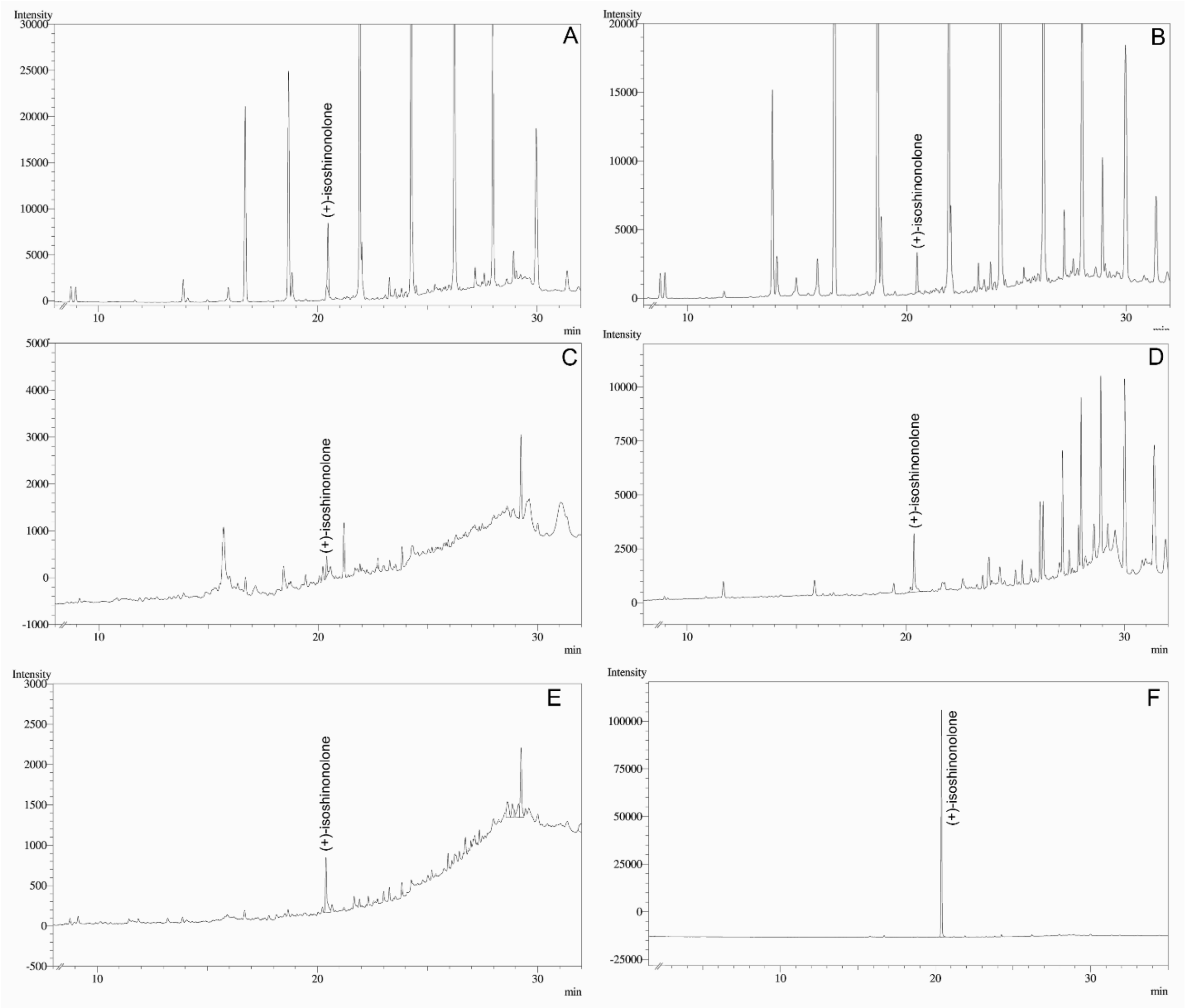


**Fig. S10. Detection of (+)-isoshinanolone in *Nepenthes* species/hybrid peristome and lid EFNs.** EFNs of **A**. *N. khasiana* peristome; **B**. *N. khasiana* lid; **C**. *N. mirabilis* peristome; **D**. *N. mirabilis* x *N. khasiana* peristome; **E**. *N. mirabilis* x *N. rafflesiana* peristome; **F**. (+)-isoshinanolone standard.

*Nepenthes khasiana*, *N. mirabilis*, *N. mirabilis* × *N. khasiana*, *N. mirabilis* × *N. rafflesiana* peristome and lid EFNs (each) were dissolved separately in 10 ml water, and partitioned with chloroform (5 × 10 ml) to obtain the peristome and lid chloroform fractions. Stock solution (1 mg/1.5 ml) of (+)-isoshinanolone was prepared in methanol and injected on to a GC-2010 Plus Gas Chromatograph with AOC-20i autoinjector and FID (Shimadzu, Japan), fitted with an Rxi-5 Sil MS capillary column (5% diphenyl/95% dimethyl polysiloxane, non-polar, 30 m x 0.25 mm i.d., 0.25 µm film thickness; Restek, USA). GC operation conditions: injection mode split; split ratio 5; injector temperature 220°C; oven temperature programme 60-150°C (5°C/min); 150-250°C (10°C/min); carrier gas $N_2$ at 3 ml/min; detector temperature 250°C.

Chloroform fractions of *Nepenthes* species/hybrid peristome and lid EFNs were made up to 1.5 ml separately in chloroform, and GC procedures were repeated to detect (+)-isoshinanolone in these fractions.

**Supplementary videos**

**Video S1.** Biotest: Behavioral pattern of *N. khasiana* peristome EFN-fed *A. gracilipes* ants.

**Video S2.** Biotest: Behavioral pattern of sugar mix + (+)-isoshinanolone-fed *A. gracilipes* ants.

**Video S3.** Biotest: Behavioral pattern of sugar mix (only)-fed *A. gracilipes* ants.

**Video S4.** Biotest: Behavioral pattern of control (water only-fed) *A. gracilipes* ants.

**Video S5.** Behavioral pattern of *A. gracilipes* ants on the peristome and leaves/tendrils of *N. khasiana*.

(Black spots seen in Videos S1, S2 and S3 are the soil/debris adhered to the ants when they were transferred to the experimental jars from their natural habitats).

(Video S5 is a combined file of four short videos, taken from the Institute Conservatory in November 2024, displaying multiple scenarios of interaction of the ant, *A. gracilipes*, with *N. khasiana*).

In biotests (24 h, *in vivo*), major symptoms observed in *N. khasiana* peristome EFN (1000 µg/ml)-, sugar mix + (+)-isoshinanolone (10 µg/ml)- and sugar mix + donepezil (10 µg/ml)-fed *A. gracilipes* ants were slow movements or cramps (muscular weakness), enhanced grooming (head, body) activity, fallen upside down, spasms and even deaths (in (+)-isoshinanolone-fed ants, 10%) (Videos S1, S2).

Ants in all three treatment groups after 24 h displayed strong AChE inhibition (*N. khasiana* peristome EFN (1000 µg/ml) 95.7 ± 1.5%, sugar mix + (+)-isoshinanolone (10 µg/ml) 97.7 ± 2.5%, sugar mix + donepezil (10 µg/ml) 91.7 ± 5.0%), whereas sugar mix-fed (10.7 ± 1.2%; Video S3) and control (unfed) (6.3 ± 0.5%; Video S4) *A. gracilipes* ants showed only negligible activity.

*A. gracilipes* ants on the peristomes of *N. khasiana* in the Conservatory showed behavioral patterns similar to test ants (Videos S1, S2), whereas (unfed) ants on its leaves and tendrils showed normal behavior (fast movements) (Video S5).
